# S1-Domain RNA-Binding Protein (CvfD) Is a New Post-Transcriptional Regulator That Mediates Cold Sensitivity, Phosphate Transport, and Virulence in *Streptococcus pneumoniae* D39

**DOI:** 10.1101/2020.06.23.167635

**Authors:** Dhriti Sinha, Jiaqi J. Zheng, Ho-Ching Tiffany Tsui, John D. Richardson, Nicholas R. De Lay, Malcolm E. Winkler

## Abstract

Post-transcriptional gene regulation often involves RNA-binding proteins that modulate mRNA translation and/or stability either directly through protein-RNA interactions or indirectly by facilitating the annealing of small regulatory RNAs (sRNAs). The human pathogen *Streptococcus pneumoniae* D39 (pneumococcus) does not encode homologs to RNA-binding proteins known to be involved in promoting sRNA stability and function, such as Hfq or ProQ, even though it contains genes for at least 112 sRNAs. However, the pneumococcal genome contains genes for other RNA-binding proteins, including at least six S1-domain proteins; ribosomal protein S1 (*rpsA*), polynucleotide phosphorylase *(pnpA*), RNase R (*rnr*), and three proteins of unknown functions. Here, we characterize the function of one of these conserved, yet uncharacterized S1-domain proteins, SPD_1366, which we have renamed CvfD (<u>C</u>onserved <u>v</u>irulence <u>f</u>actor <u>D</u>), since loss of this protein results in an attenuation of virulence in a murine pneumonia model. We report that deletion of *cvfD* impacts expression of 144 transcripts including the *pst1* operon, encoding the phosphate transport system 1 in *S. pneumoniae*. We further show that CvfD post-transcriptionally regulates the PhoU2 master regulator of the pneumococcal dual phosphate transport system by binding *phoU2* mRNA and impacting PhoU2 translation. CvfD not only controls expression of phosphate transporter genes, but also functions as a pleiotropic regulator that impacts cold sensitivity and the expression of sRNAs and genes involved in diverse cellular functions, including manganese uptake and zinc efflux. Together, our data show that CvfD exerts a broad impact on pneumococcal physiology and virulence, partly by post-transcriptional gene regulation.

**SIGNIFICANCE:** Recent advances have led to the identification of numerous sRNAs in the major human respiratory pathogen, *S. pneumoniae*. However, little is known about the functions of most sRNAs or RNA-binding proteins involved in RNA biology in pneumococcus. In this paper, we characterize the phenotypes and one target of the S1-domain RNA-binding protein CvfD, a homolog of “general-stress protein 13” identified, but not extensively characterized in other *Firmicute* species. Pneumococcal CvfD is a broadly pleiotropic regulator, whose absence results in misregulation of divalent cation homeostasis, reduced translation of the PhoU2 master regulator of phosphate uptake, altered metabolism and sRNA amounts, cold sensitivity, and attenuation of virulence. These findings underscore the critical roles of RNA biology in pneumococcal physiology and virulence.

## INTRODUCTION

Over the past decade there has been increasing evidence regarding the importance of post-transcriptional gene regulation in modulating the physiology and virulence of Gram-positive bacterial pathogens (1–3). Post-transcriptional regulation of gene expression is often carried out by RNA-binding proteins that frequently act in tandem with small regulatory RNAs (sRNAs) (4). Some RNA-binding proteins bind mRNAs altering their transcription, translation, or stability (5, 6). In many cases, the activity of these RNA-binding proteins is modulated by sRNAs, which upon expression can act as antagonists, blocking protein binding to target mRNAs (7). For example, in the Gram-positive bacterial pathogens *Enterococcus faecalis* and *Listeria monocytogenes,* the two-component system (TCS) response regulator EutV upon phosphorylation binds to a dual hairpin in the 5’ untranslated region (5’-UTR) of the polycistronic mRNA encoding ethanolamine utilization genes. This binding stabilizes an anti-terminator that promotes transcription elongation (8, 9). EutV function is antagonized by the EutX/Rli55 sRNAs in the presence of ethanolamine and the absence of B_12_, blocking it from binding the *eut* mRNA. As a consequence, a terminator forms in the 5’-UTR of the *eut* mRNA, resulting in transcriptional attenuation (8, 9). In *Bacillus subtilis* and *Escherichia coli,* the RNA-binding protein CsrA binds to mRNAs causing translational repression (7, 10, 11). In *E. coli*, CsrA is sequestered by at least two sRNAs, CsrB and CsrC (12, 13), whereas in *B. subtilis* CsrA is antagonized by the protein FliW, which inhibits its RNA-binding activity through binding to an allosteric site on CsrA (14).

Other RNA-binding proteins work in concert with sRNAs that act by base pairing with their respective mRNA targets, altering their transcription (15), translation (16), or stability (17). sRNAs are a well-studied class of post-transcriptional regulators that have been implicated in controlling a wide variety of physiological responses in bacteria ranging from stress responses to virulence gene expression (4, 18, 19). One bacterial RNA-binding protein that has gained much attention is the RNA chaperone protein Hfq (20–23). Hfq stabilizes sRNAs by binding and protecting them from ribonucleolytic cleavage (24), but also facilitates annealing between sRNAs and their target mRNAs (25–28). In Gram-negative pathogens including *Salmonella enterica* (29, 30), *Yersinia pestis* (31, 32), and *Vibrio cholerae* (33, 34), deletion of *hfq* results in pleiotropic phenotypes affecting growth, the ability to cope with environmental stresses, and virulence properties. In the Gram-positive pathogen *Clostridium difficile* depletion of *hfq* induces multiple phenotypic changes including altered growth, cell morphology, stress response, sRNA abundance, and biofilm formation (35). Virulence phenotypes have also been reported for an *hfq* deletion mutant in *Staphylococcus aureus* (36, 37) and *Listeria monocytogenes* (38, 39); however, in contrast to Gram-negative pathogens, the role of Hfq in RNA regulation in Gram-positive pathogens is far less established (40–42). Moreover, important Gram-positive pathogens, such as *Streptococci* and *Mycobacteria*, lack a recognizable Hfq homolog.

*S. pneumoniae* is a Gram-positive, low GC-content aerotolerant anaerobe that exclusively resides within the human respiratory tract as a commensal bacterium, but can become an opportunistic pathogen (43). *S. pneumoniae* is the leading cause of bacterial pneumonia and other serious infections, including otitis media, sinusitis, meningitis, and septicemia, and over 1 million people succumb to pneumococcal infections annually worldwide (44, 45). Prior studies have reported the presence of over one hundred sRNAs both in the serotype 2 (D39) and serotype 4 (TIGR4) strains of *S. pneumoniae*, using a combination of computational predictions, high-throughput RNA sequencing, and northern blot analyses (46–51). Interestingly, a recent study by Zheng et al. (52) identified the KH-RNA-binding domain proteins, KhpA and KhpB, as post-transcriptional regulators of the cell division protein FtsA in *S. pneumoniae*; however, the mechanism underlying this regulation remains unresolved. More recently, hundreds of potential RNA-protein complexes were identified in the TIGR4 strain of *S. pneumoniae* using a Grad-seq based approach, and further analysis of those interactions uncovered a new function for the exonuclease Cbf1 in stabilizing certain specific pneumococcal sRNAs by trimming their 3’ end, which presumably removes a binding site for other RNases (53). There has been no other reported evidence for any RNA-binding protein that may function similar to Hfq in *S. pneumoniae* or other pathogenic *Streptococci*. The structure of the pneumococcal RNA-binding protein CvfB (<u>C</u>onserved <u>v</u>irulence <u>f</u>actor B) was solved, revealing the presence of 3 consecutive S1 RNA-binding domains and a fourth unique WH (winged helix) RNA-binding domain (54). The S1(3)-WH domains have been further shown to possess RNA-binding properties with the following preference for polyribonucleotides – poly(U)> poly(A)> poly(C)> poly(G) (54). Although little is known about the roles of S1-domain proteins in RNA-mediated gene regulation in Gram-positive bacteria, S1-domain containing proteins are of particular interest because of some of their functional similarities with Hfq (55).

In Gram-negative bacteria, Hfq and ribosomal protein S1 consist of six modules that preferentially bind A, U, or A/U-rich sequences and are reported to impact transcription, translation, and RNA degradation (55). S1 domains fall under the OB (<u>o</u>ligonucleotide/oligosaccharide <u>b</u>inding)-fold superfamily which also contain the related RNA-binding cold-shock domains (56, 57). S1-domains use the common OB-fold binding surface as the RNA-recognition core, which in turn, are comprised of two β-strands and several surrounding loops. The protein-RNA complex is formed via stacking interactions between the nucleic acid bases and conserved aromatic residues on the two-stranded β-sheet core, and these interactions are further stabilized by surrounding loops and secondary structure elements. The RNA-binding face consists of five highly conserved residues that are located in the turns between β-strands 1 and 2 (Phe-19), in the middle of strand 2 (Phe-22), at the end of the strand 3 (His-34), and in the turn between the final two strands of sheet 5 (Asp-64, Arg-68) (58). In 3-D structures, all of these residues are close to one another and are predicted to be on the surface of one side of the antiparallel β-barrel (56–59). Interestingly, studies have also shown that S1-domains can bind pseudoknot structures and U or A/U-rich mRNA leader regions upstream of Shine-Delgarno sequences with high affinities (55).

Genome-wide BLAST searches revealed the presence of several conserved-S1-domain containing RNA-binding proteins in *S. pneumoniae* strain D39 that included CvfB, the exoribonucleases PNPase (polynucleotide phosphorylase) and RNase R, the endoribonuclease YbeY, and SPD_1366, a protein of unknown function. SPD_1366 is alternatively called general stress protein 13 (GSP13) or YugI in *B. subtilis* (60) and Ygs in *Staphylococcus epidermidis* (61). YugI homologs are present only in Gram-positive bacteria and possess a single conserved N-terminal S1-domain and a unique C-terminal charge-rich tail (Figure 1). In both *B. subtilis* and *S. epidermidis*, this protein is induced under certain stress conditions like ethanol stress, heat shock, cold shock and salt stress (60, 61). In *B. subtilis*, the protein can also be induced under glucose starvation, oxidative stress, and ammonium starvation (60). Previous studies have also shown that YugI can associate with ribosomes during growth in exponential phase (60). However, its function in ribosomes, if any, remains unknown. On the other hand, in *S. epidermidis*, Ygs was shown to impact biofilm formation and biofilm-mediated infections (61). Whether YugI or Ygs can function as potential RNA-binding proteins in the above-mentioned Gram-positive bacteria remains an open question.

**Figure 1:**
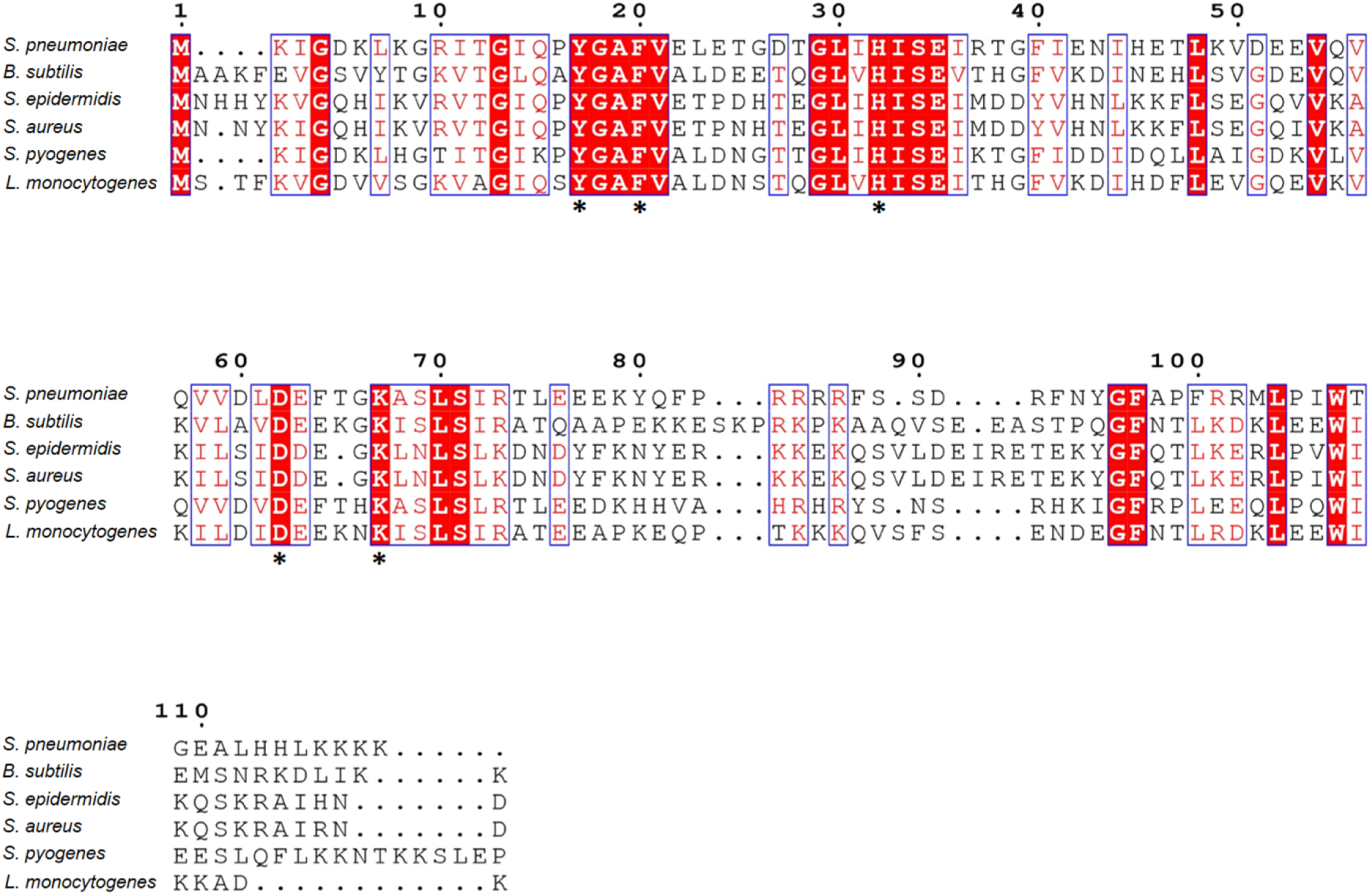
Multiple sequence alignment of CvfD homologs from several Gram-positive bacteria. Alignment of the CvfD homologs from *S. pneumoniae* (sequence ID ABJ54781.1), *B. subtilis* (sequence ID AGG62546.1), *S. epidermidis* (sequence ID OXE79056.1), *S. aureus* (sequence ID KMR00465.1), *S. pyogenes* (sequence ID SQG97127.1), and *L. monocytogenes* (sequence ID NP_465892.1) was obtained by using ClustalOmega. The residues marked by (*) indicate conserved RNA-binding residues in the S1-domain as predicted by Bycroft et al., (1997). Residues in the S1-domain that are conserved among all of the different Gram-positive bacteria listed here are shaded in red.

In this study, we investigated the function of S1-domain protein SPD_1366 in *S. pneumoniae* serotype 2 strain D39 by studying the phenotypes exhibited by deletion mutants of *spd_1366*. We renamed SPD_1366 as <u>C</u>onserved <u>V</u>irulence <u>F</u>actor <u>D</u> (CvfD) based on its attenuation of pneumococcal virulence *in vivo*. We also show that CvfD is an RNA-binding protein regulator that controls the expression of the PhoU2 master regulator of the dual phosphate transport system at the post-transcriptional level. We demonstrate that deletion of *cvfD* leads to cold sensitivity and numerous changes in relative mRNA and sRNA transcript amounts, some of which are reflected by Δ*cvfD* phenotypes, such as misregulation of divalent cation homeostasis. Altogether, these results support the conclusion that CvfD is a highly pleiotropic post-transcriptional regulator in *S. pneumoniae*.

## RESULTS

### CvfD is required for growth of *S. pneumoniae* D39 at both optimal (37°C) and lower (32°C) temperatures

CvfD, a 119 amino-acid GSP13-like protein, is one of six S1 RNA-binding-domain proteins encoded in the genome of *S. pneumoniae* strain D39. The S1 domain of CvfD spans residues 2 to 73 (Fig. 1). CvfD shares 40% and 44% identity at the amino acid level with *B. subtilis* YugI and *S. epidermidis* Ygs, respectively. Since prior studies implicated the CvfD homologs YugI and Ygs in stress adaptation (60, 61), we constructed an in-frame markerless clean-deletion or insertional-deletion in *cvfD* (Table S1 and S2) to test for its impact on pneumococcal growth and physiology. We found that CvfD is required for growth of *S. pneumoniae* D39 at the optimal temperature of 37°C in aged BHI broth, as the Δ*cvfD* mutant exhibited a significant defect in growth rate and growth yield when compared to the wild-type (WT) parent strain (Fig. 2A, S1A and S1B; Table S3). These growth defects were more pronounced in BHI medium that remained in sealed bottles at room temperature for two weeks or longer than in freshly prepared medium (Fig. S1A and S1B), indicating some change in this medium that affects the Δ*cvfD* mutant, but not the WT strain. In 15-24-day-old BHI broth, designated as “aged” and used in most experiments in this paper, the average doubling time and growth yield for the Δ*cvfD* mutant were ≈58 min and ≈0.3, respectively, compared with ≈45 min and ≈0.9 for the WT strain, respectively. By contrast, *cvfD* insertions did not appreciably affect the fitness for growth of the serotype 4 TIGR4 strain in a minimal medium containing several different carbon sources in a Tn-Seq screen (45). Likewise, *ygs* mutants of *S. epidermidis* defective in the pneumococcal *cvfD* homolog did not strongly affect growth (61).

**Figure 2:**
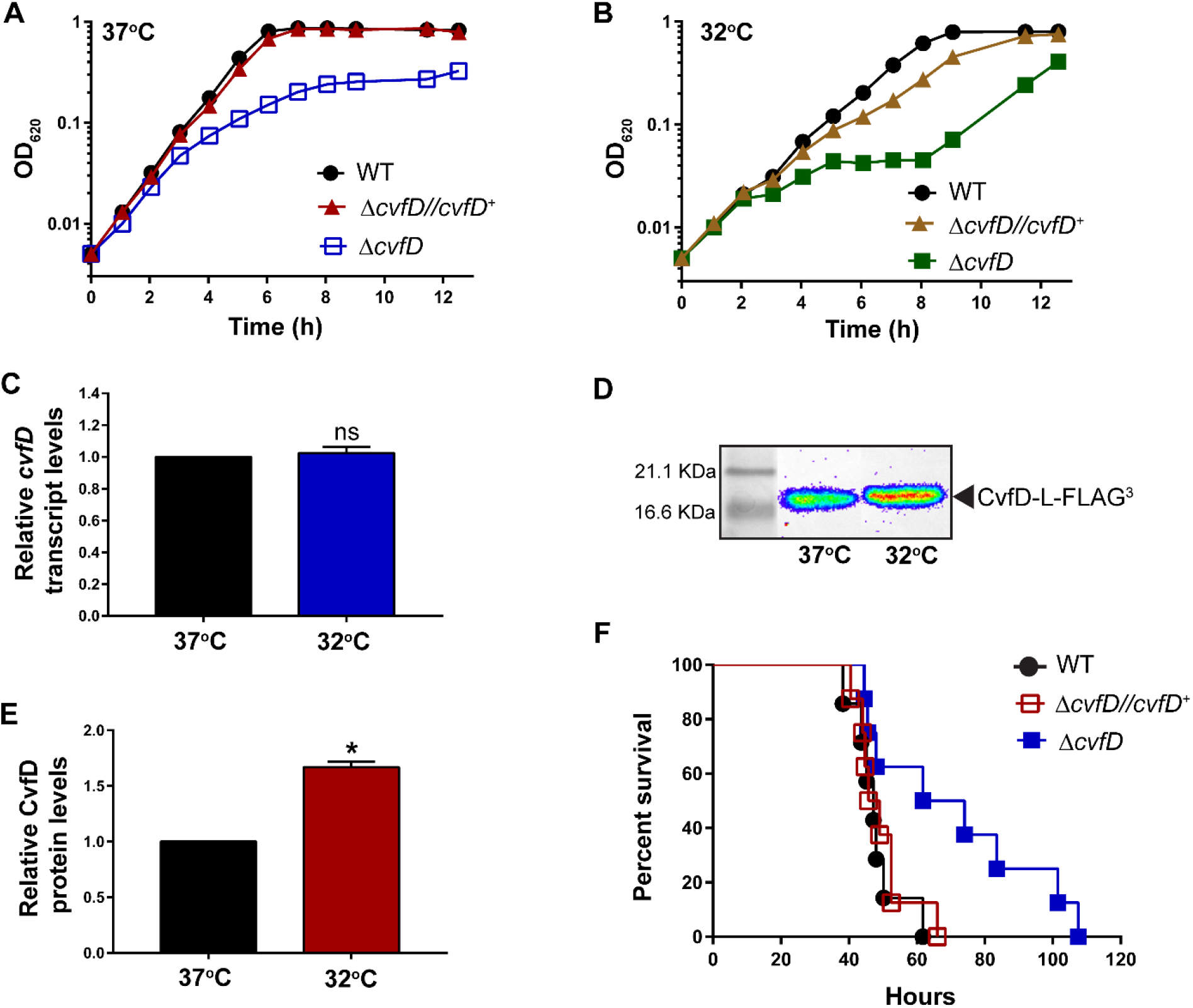
Growth and virulence phenotypes of Δ*cvfD* mutant. (A, B) Growth characteristics of encapsulated D39 parent strain (IU1781) and its derived mutant strains, Δ*cvfD* (IU4772) and Δ*cvfD bgaA*::P_*ftsA*_ *cvfD*^+^ (IU5508), at 37°C and at 32°C in aged (24-day old) BHI broth. The accelerated growth rate of the Δ*cvfD* (IU4772) mutant at 32°C after 8 hours of incubation is due to accumulation of spontaneous suppressor mutations. At least five independent growth curves were performed at 37°C or 32°C in aged BHI with similar results, and a representative curve is shown. Average growth rates and growth yields are listed in Supplemental Table S3. (C) qRT-PCR analysis to determine *cvfD* transcript steady state levels in the parent strain (IU1781) relative to *gyrA* (internal control) at 37°C and 32°C. Primers used for qRT-PCR analyses are listed in Supplemental Table S5. (D, E) Western blot analysis to determine CvfD protein levels at 37°C and 32°C in an isogenic strain expressing CvfD-L-FLAG^3^ (IU8717). (C, D, E) All strains were grown to early exponential phase in aged BHI broth and data points and error bars represent the means and the standard errors (SEM) of at least three independent experiments. Asterisks (*) and ns indicate P<0.05 and not significantly different, respectively. (F) Survival curve analysis showing disease progression in an invasive model of pneumonia. ICR male mice were inoculated intranasally with ~10^7^ CFUs in 50 μL inoculum of either the D39 parent (IU1781) strain or its derived isogenic mutants (Δ*cvfD*; IU4772 or Δ*cvfD bgaA*::P_*ftsA*_ *cvfD*^+^; IU5508 (complemented strain)). Eight animals were infected per strain and disease progression was followed in real time by a survival curve analysis (*Materials and Methods*). Survival curves were analyzed by Kaplan-Meier statistics and log-rank tests to determine P-values.

S1-domain proteins often mediate bacterial cold-shock responses (59, 60). Therefore, we characterized the growth of a Δ*cvfD* mutant strain at a lower growth temperature (32°C) to test for a cold-sensitivity (CS) phenotype. The Δ*cvfD* mutant had a considerably lower growth rate and yield when grown at 32°C compared to 37°C in aged BHI broth (Fig. 2B, S1C, and S1D; Table S3), indicating a CS phenotype. Again, the relative growth rate of the Δ*cvfD* mutant was much slower in aged BHI broth than in fresh BHI broth (Fig. S1D), although the relative growth yield was markedly lower at 32°C for the Δ*cvfD* mutant in all batches of BHI broth (Fig. S1C). Prolonged incubation of Δ*cvfD* cultures at 32°C led to the accumulation of faster growing suppressor mutations (Fig. 2B), which are analyzed below. To further confirm the CS phenotype of the Δ*cvfD* mutant, the Δ*cvfD* mutant and WT were streaked onto a different medium (TSAII-blood) in agar plates, and colony size was observed. At 37°C, the colonies of the Δ*cvfD* mutant were only marginally smaller than those of the WT strain (Fig. S2A), whereas the Δ*cvfD* mutant did not grow at all at 32°C (Fig. S2B). Finally, the growth defects of the Δ*cvfD* mutant in aged BHI broth or at 32°C were complemented by expressing the WT *cvfD*^+^ gene from a constitutive P_*fts*A_ promoter at the ectopic *bgaA* locus (Fig. 2A, 2B, S2A, and S2B; Tables S2 and S3) (49). We conclude that Δ*cvfD* mutant growth is defective at 37°C in some media conditions and that the *cvfD*^+^ gene is required for growth at 32°C, near the temperature in the nasopharynx of humans (62).

### CvfD is post-transcriptionally induced under cold stress in *S. pneumoniae*

The CS phenotype of the Δ*cvfD* mutant prompted us to investigate whether CvfD can function as a cold-shock protein in *S. pneumoniae* D39. Studies in *E. coli* show that the S1 domain from the exoribonuclease PNPase complements the CS phenotype of a Δ*cspA* mutant; CspA is one of several well-studied CSPs in *E. coli* (63). The PNPase S1-domain and the CvfD homolog S1-domain from *B. subtilis* consist of the same conserved RNA-binding residues and have the same overall protein structure (60). Furthermore, sequence alignments of the *B. subtilis* CvfD homolog YugI with *S. pneumoniae* CvfD reveals conservation of the same RNA-binding residues (Fig. 1). Expression of CSPs, such as CspA from *E. coli,* is induced under cold stress due to increased transcription, mRNA stabilization, and translation (64). We tested whether the lower temperature of 32°C, which mimics cold stress like conditions in *S. pneumoniae*, can result in an increased amount of the *cvfD* mRNA and/or CvfD protein. Quantitative real-time PCR (qRT-PCR) show that the relative mRNA level of *cvfD* did not significantly increase at 32°C compared to 37°C (Fig. 2C). Next, we determined the relative amount of CvfD at low and optimal growth temperatures using a C-terminal linker-FLAG^3^ (L-FLAG^3^) epitope-tagged construct of CvfD expressed from its native locus. Introduction of the CvfD-L-FLAG^3^ tagged-protein did not cause any obvious growth defect in contrast to a Δ*cvfD* mutant (data not shown). We did observe a modest, but significant increase in CvfD protein levels (≈1.6 fold) at 32°C compared to 37°C, as determined by quantitative western blotting (Fig. 2D and 2E). The observation that relative CvfD protein levels increase without a corresponding increase in *cvfD* transcript amount suggests that CvfD expression is controlled at the post-transcriptional level, potentially by an autoregulatory mechanism. Taken together, our data are consistent with the notion that CvfD acts as a CS protein in *S. pneumoniae* D39.

### CvfD is important for pneumococcal pathogenesis

Prior studies in *S. epidermidis* show that the CvfD homolog Ygs is required for biofilm-associated infections in a catheter-mouse model (61). In addition, Tn-seq studies in serotype 4 strain (TIGR4) of *S. pneumoniae* suggested that loss of the CvfD homolog (SP_1537) leads to a significant defect in lung colonization in a murine model of infection (45). To verify this virulence attenuation, we tested the impact of the D39 Δ*cvfD* clean-deletion mutation on pneumococcal disease progression relative to a WT parent (*cvfD*^+^) strain in a murine model of invasive pneumonia. Mice were infected with an intranasal dose of ≈10^7^ CFUs in 50 μL and subsequent disease progression was monitored by checking survival curve analysis in which death was not used as an endpoint (see *Materials and Methods*). Mice infected with Δ*cvfD* mutant were substantially attenuated for virulence with a median survival time of 68 h (P_val_ = 0.03), which was significantly longer than the 47 h median survival time of mice inoculated with the WT *cvfD*^+^ parent (Fig. 2F). Finally, the Δ*cvfD* strain complemented with *cvfD*^+^ constitutively expressed from an ectopic locus caused a median survival time (47 h) similar to that of the WT strain, but significantly different (P_val_ = 0.04) from that displayed by the Δ*cvfD* strain. Thus, ectopic expression of CvfD completely complemented the attenuation of the Δ*cvfD* mutant. Altogether, these results indicate that CvfD is a virulence factor in *S. pneumoniae* D39.

### CvfD regulates the relative amounts of numerous mRNA transcripts

To gain insight into the mechanism by which CvfD influences pneumococcal growth and virulence, we investigated its impact on gene expression in *S. pneumoniae* D39. We compared the genome-wide transcriptome expression profiles of a Δ*cvfD* mutant (IU5506) with that of its isogenic WT parent strain (IU3116). Strains were grown in exponential phase to an OD_620_ ≈0.1 in matched batches of aged BHI broth at 37°C in an atmosphere of 5% CO_2_, and RNA-seq analysis was performed as described in *Materials and Methods*. mRNA-seq data analysis revealed that 124 transcripts were significantly differentially expressed above a fold change cut-off of 1.8 and a P_adj_-value cut-off of 0.005. Transcripts from 78 or 46 open reading frames (ORFs) were up- or down-regulated, respectively, in the Δ*cvfD* mutant compared to the WT strain (Table 1, Fig. 3A). The largest fraction of up-regulated genes mediate phosphate ion uptake, transport of metal ions, carbohydrate metabolism, energy production, and virulence (Table 1). In particular, relative transcript abundance for genes encoding the phosphate transport system 1 (*pst1*) operon (*pstSCAB*) and its regulator *phoU1* increased by ≈4 fold in the Δ*cvfD* strain, whereas the relative amounts of transcripts encoding the *psaBCA* manganese transporter and the zinc efflux pump (*czcD*) increased by ≈2 and ≈4 fold, respectively. The relative expression of transcripts corresponding to important enzymes in glycolysis (*gap* and *pgk*) were up-regulated by 2-3 fold in the Δ*cvfD* mutant. Notably, lack of CvfD resulted in a 2-3 fold increase in the relative expression of genes encoding pneumococcal virulence factors (*prtA*, *pspA*, *phtA*). The largest fraction of down-regulated relative transcript amounts in the Δ*cvfD* mutant corresponded to genes encoding an iron uptake system (*piuC, piuD and piuA*; 4-5 fold), tryptophan transport (*spd_0955* and *spd_0956*; ≈2 fold) and biosynthesis (*trpABCDEFG*; 3-4 fold) genes, transfer RNAs (tRNA^Gly^, tRNA^Ile^, tRNA^Met^, tRNA^Glu^ and tRNA^Asp^; ≈2-5 fold), and genes under the control of the two-component systems (TCS) WalRK (*spd_0104* and *spd_1874*; 2-fold) and CiaRH (*htr*A and *spd_*2069; ≈3 fold), including down regulation of the *ciaRH* mRNA itself (≈2 fold).

**Table 1:**
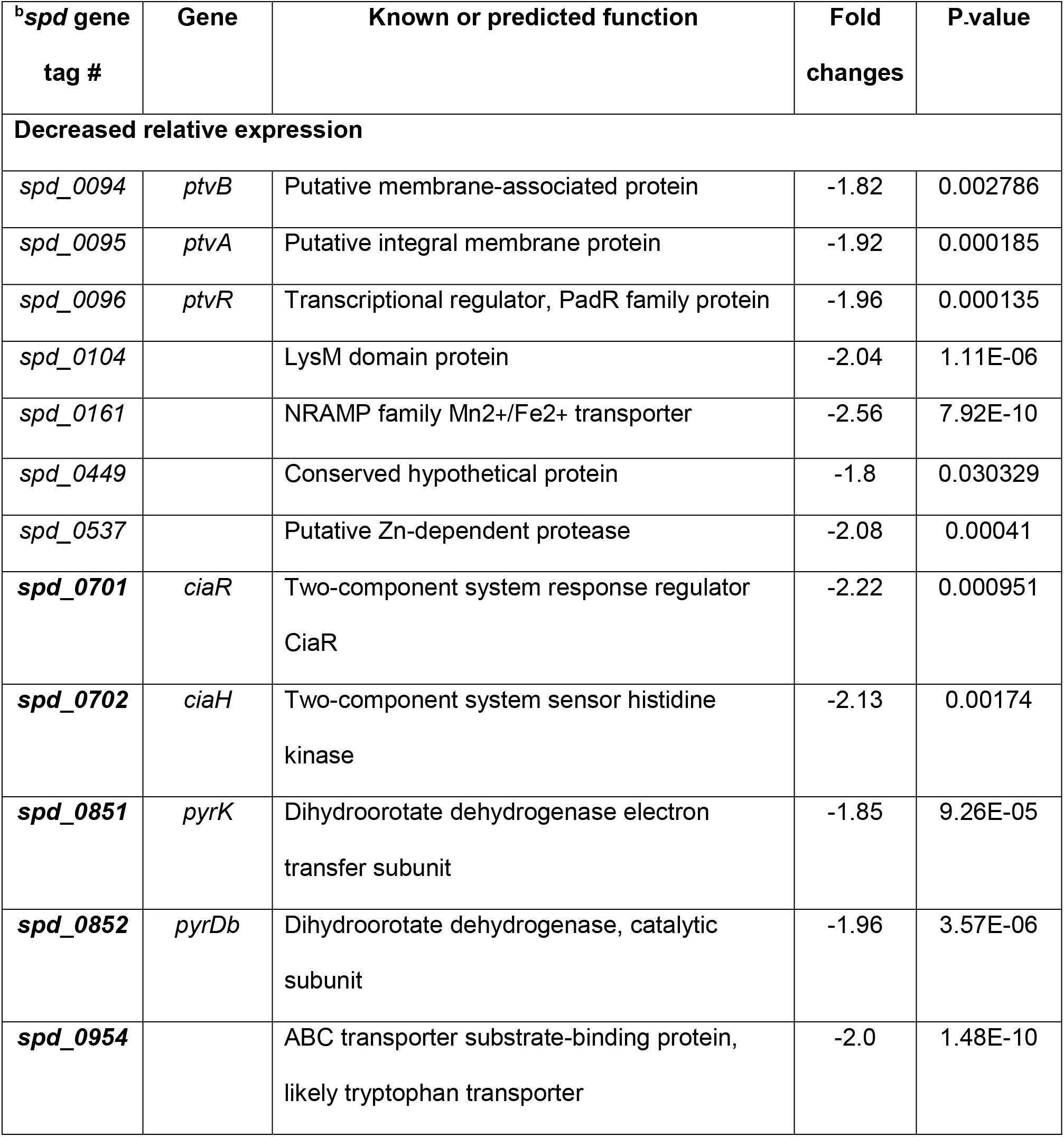
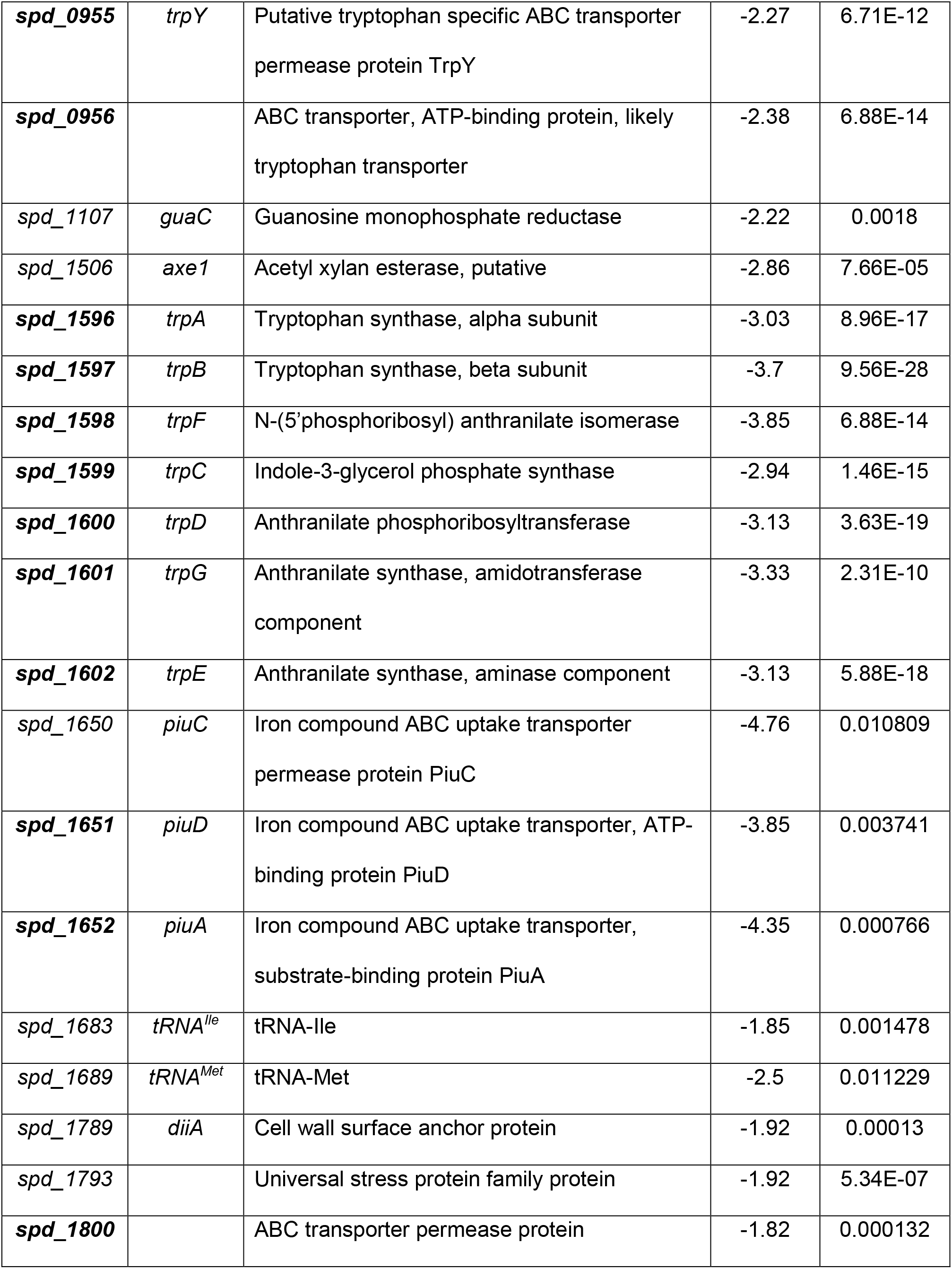
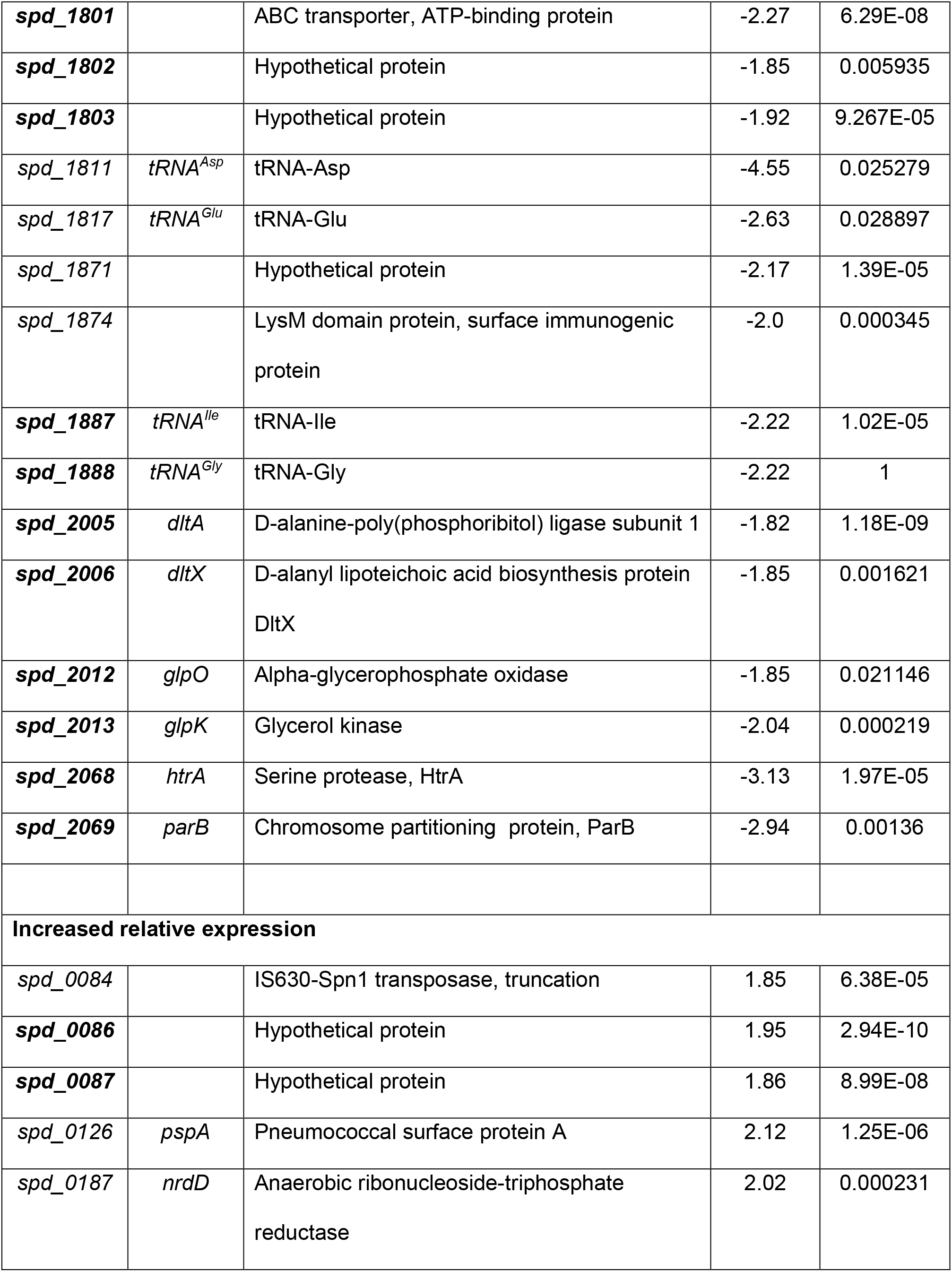
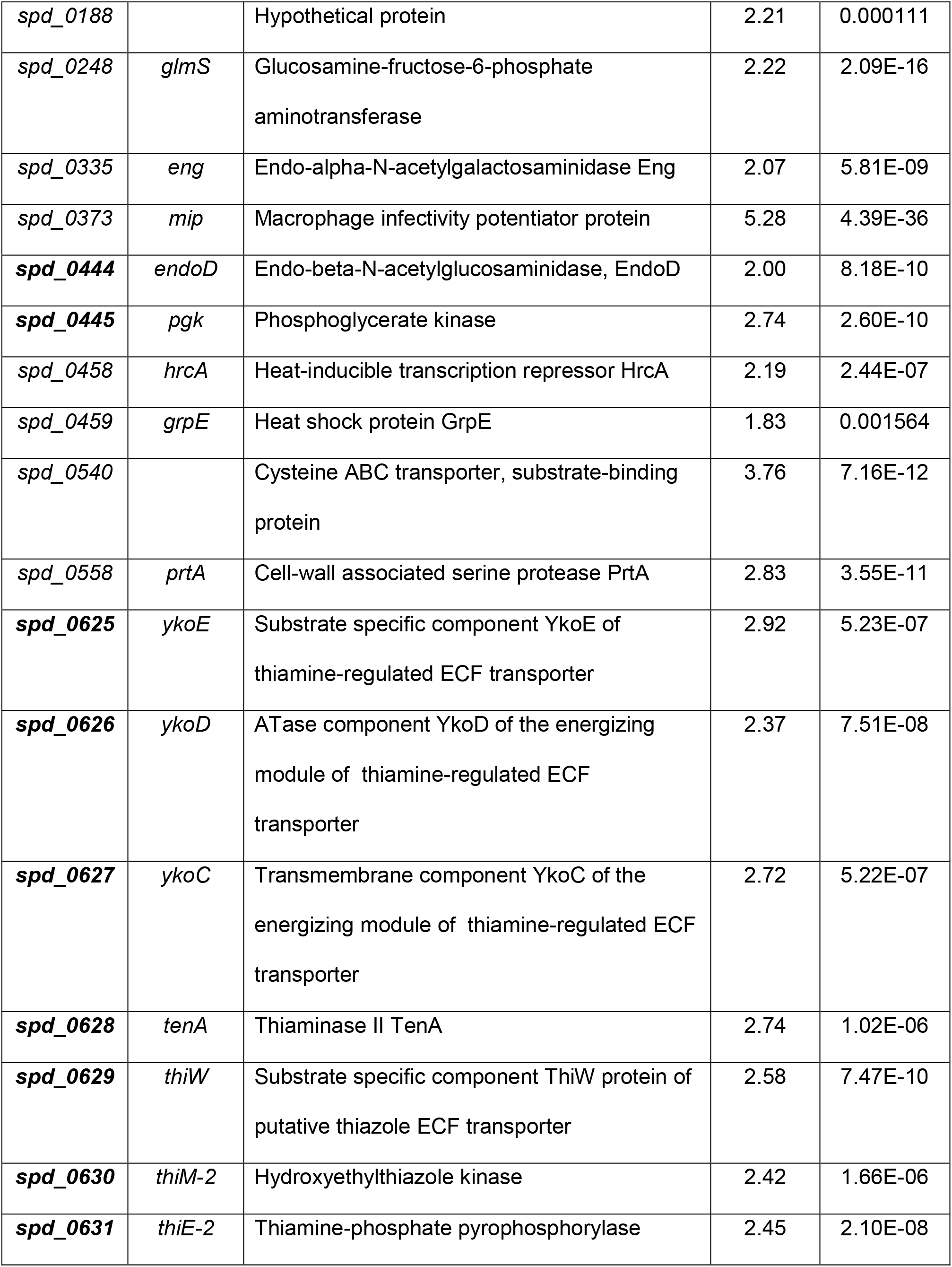
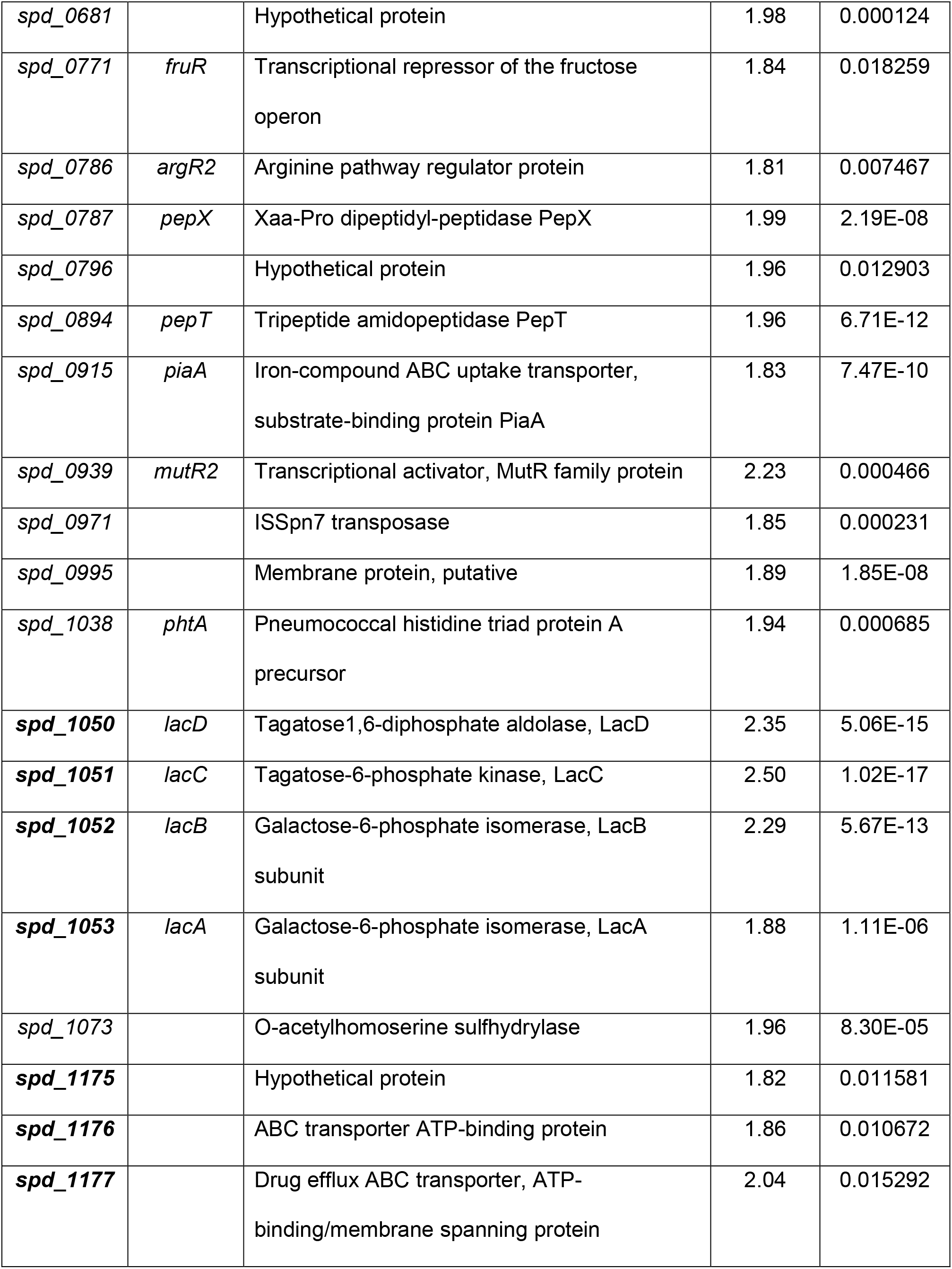
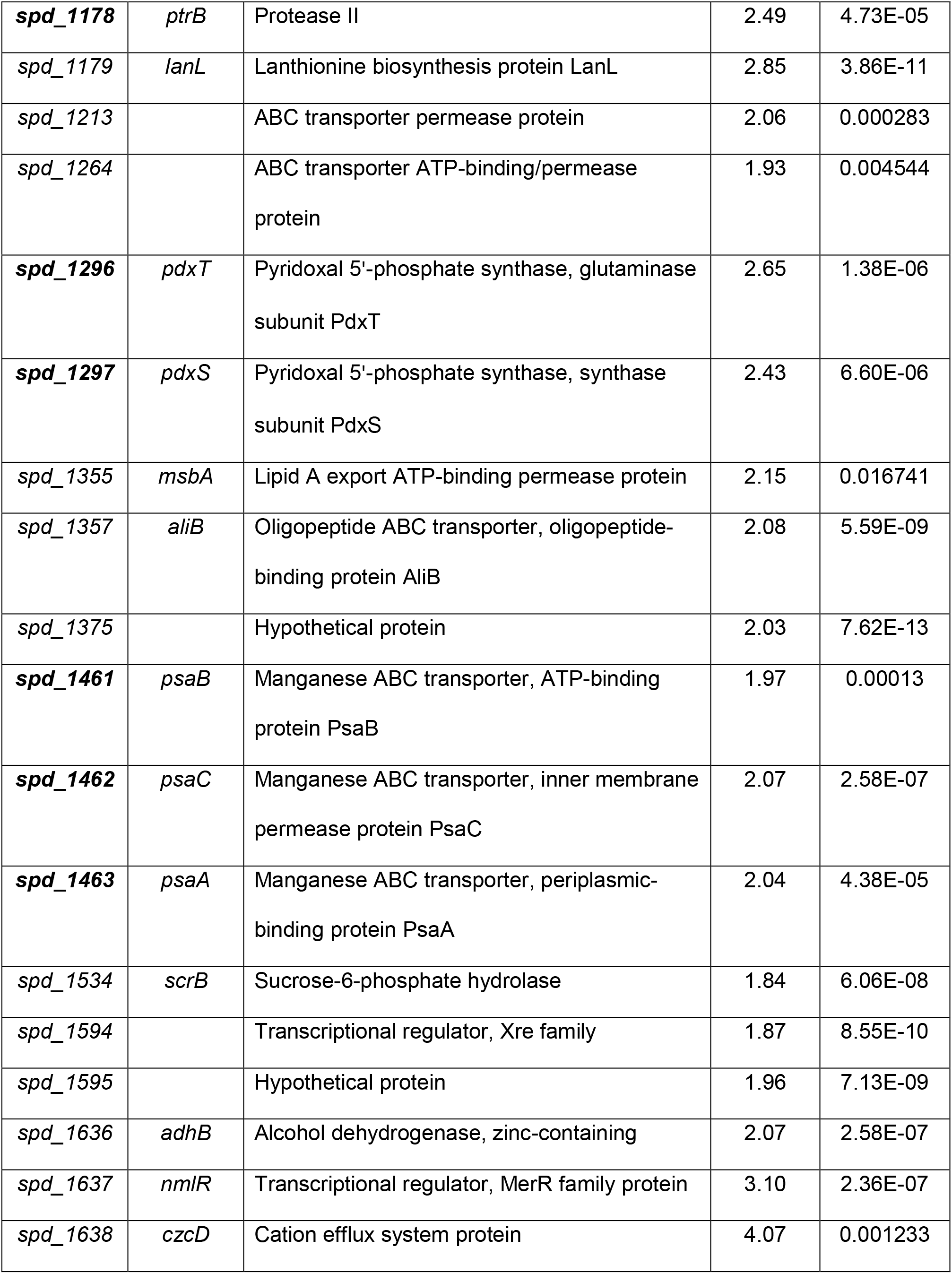
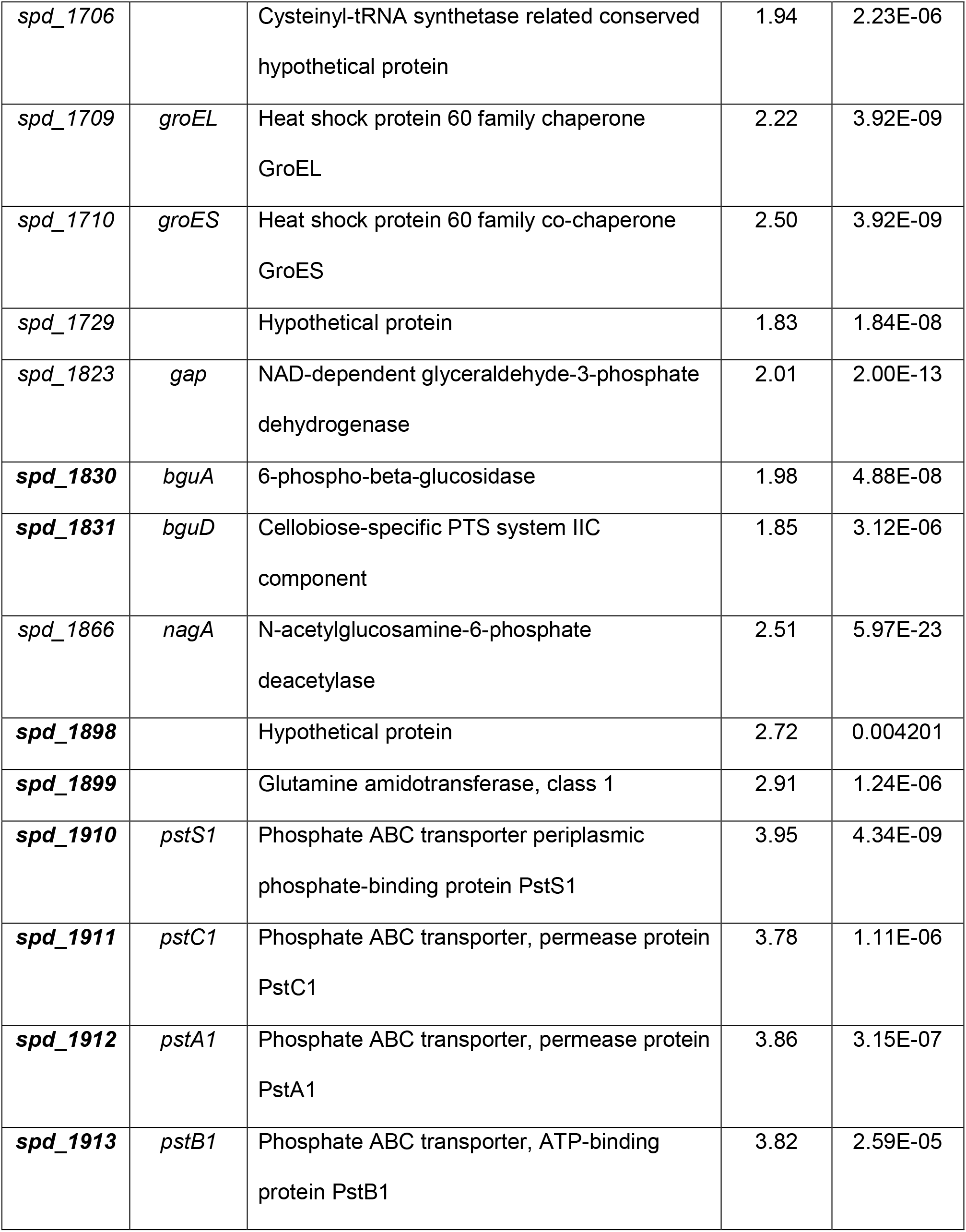
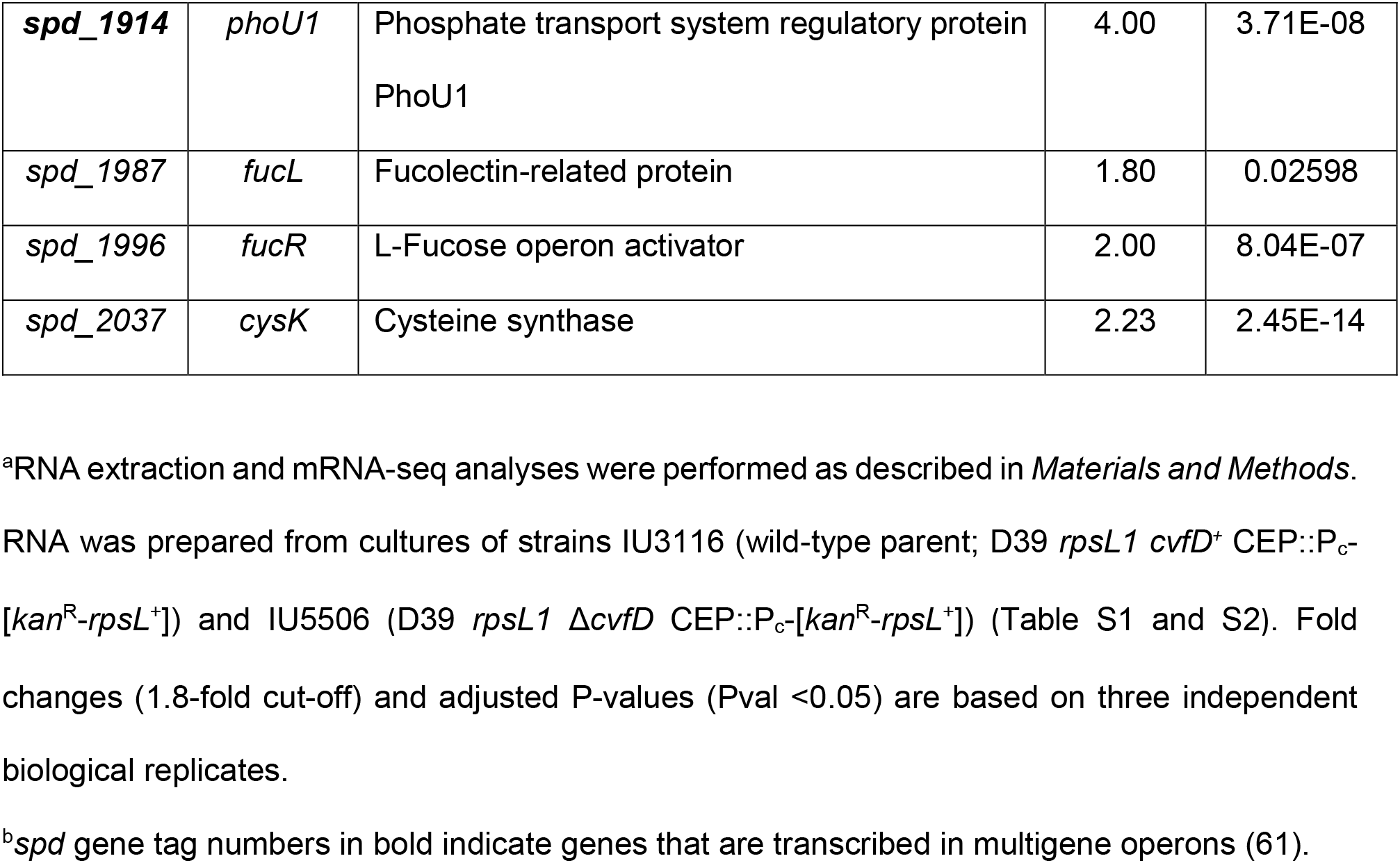
List of genes showing changes in relative transcript amounts in a Δ*cvfD* mutant compared to the *cvfD*^+^ parent strain under exponential growth in BHI broth^a^.

**Figure 3:**
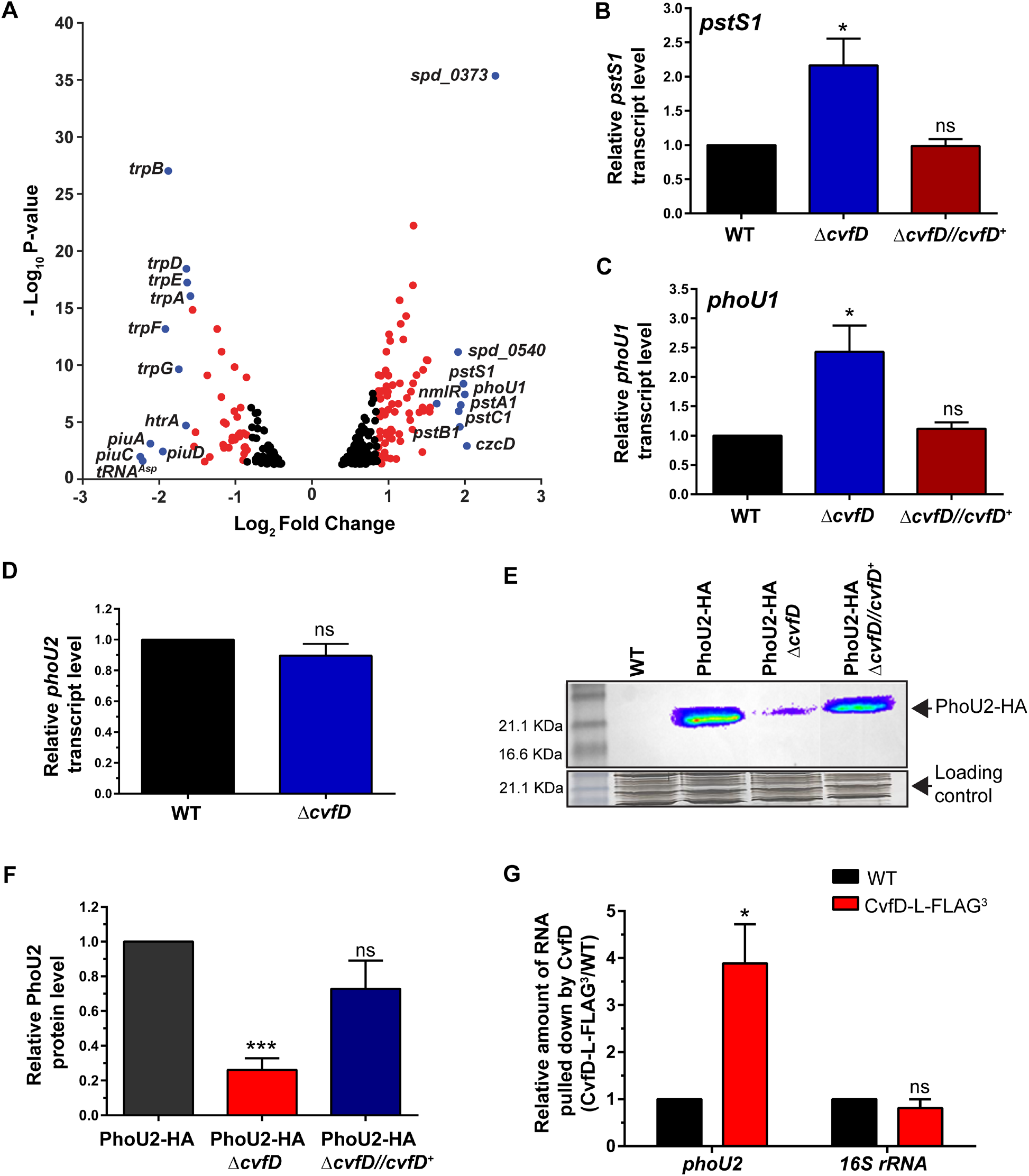
CvfD post-transcriptionally regulates the PhoU2 master regulator. (A) Volcano plot showing genome-wide changes in transcript levels from ORFs in a Δ*cvfD* mutant relative to the D39 parent strain. RNA extracted from exponential growth phase cultures of the wild-type D39 parent (IU3116; D39 *rpsL1* CEP::P_c_-[*kan*^R^-*rpsL*^+^]) and its derived isogenic Δ*cvfD* mutant (IU5506; D39 Δ*cvfD rpsL1* CEP::P_c_-[*kan*^R^-*rpsL*^+^]) in triplicate and were prepared for mRNA-seq analysis as described in *Materials and Methods*. Red and blue dots represent genes with relative transcript changes over a 1.8 fold cut-off (Log_2_fold change =0.85) and 3 fold cut-off (Log_2_fold change =1.5), respectively, with an adjusted P-value cut-off of 0.05, which are considered to be significant changes. The genes displaying changes below the above mentioned fold change and P-value cut-off are considered to be insignificant changes and are color coded in black. X-axis represent gene fold changes and Y-axis represent corresponding P-values plotted in a logarithmic scale. Genes that were significantly up-regulated or down-regulated in a Δ*cvfD* mutant compared to the parent are listed in Table 1. (B, C, D) qRT-PCR analysis was used to determine *pstS1, phoU1* and *phoU2* transcript levels in a wild-type D39 parent (WT; IU1781) and its derived isogenic mutants (Δ*cvfD*; IU4772 and Δ*cvfD bgaA*::P_*ftsA*_ *cvfD*^+^; IU5508). Transcript signal intensities were normalized to the *gyrA* transcript level, which served as the internal control, and subsequently, the expression level relative to the WT strain IU1781, which was set to 1, was calculated. (E) Western blot analysis to determine PhoU2 protein levels. Samples were prepared for western blotting from exponential phase cultures of a wild-type strain (WT; IU1781), and its derived strains *phoU2*-HA (IU8675), *phoU2*-HA Δ*cvfD* (IU8722) and *phoU2*-HA Δ*cvfD bgaA*::P_*ftsA*_ *cvfD*^+^ (IU8719) with anti-HA antibody. Representative western blots are shown in (E). Quantification of PhoU2 levels from those western blots are shown in (F), and PhoU2 expression levels are calculated relative to that in a WT strain IU1781, which was set to 1. (G) RNA-immunoprecipitation (RIP) coupled with qRT-PCR (RIP-qRT-PCR) was used to determine interactions between *phoU2* mRNA and CvfD. Strains used in RIP experiments were derived from an unencapsulated isogenic variant of D39. CvfD was immunoprecipitated from exponential phase cultures of wild-type (WT; IU1945) and an isogenic strain expressing CvfD-L-FLAG^3^ from its native locus (IU5809) using anti-FLAG antibody (Materials and Methods). RNA extracted from the elution fractions were reverse transcribed to cDNA and subsequently used as templates for qRT-PCR to determine relative levels of the *phoU2* transcript co-IPed from lysates of a strain expressing CvfD-L-FLAG^3^ compared to that of an untagged strain (WT); *phoU2* expression level was set to 1, and 16S rRNA was used as an internal control. Data shown in B, C, D, F and G represent the means (± SEM) of at least three independent experiments. Asterisks (*), (***) and ns indicate P<0.05, P<0.0001 and not significantly different, respectively. Primers used for qRT-PCR analyses are listed in Supplemental Table S5.

Results from mRNA-seq analyses were confirmed by qRT-PCR as described in *Material and Methods*. Three independent replicates of RNA samples were extracted from the following strains: WT parent (IU1781); Δ*cvfD* mutant (IU4772); and the Δ*cvfD* mutation complemented by ectopic *cvfD*^+^ expression (IU5508). Consistent with the RNA-seq results, the relative transcript levels of *pstS1*, *phoU1*, *psa*A, and *czcD* increased by 2.2, 2.6, 1.5, and 2.7 fold, respectively, in the Δ*cvfD* mutant compared to WT strain (Fig. 3B, 3C, 4A and 4B). Consistent with the RNA-seq results, the relative amounts of the *spd_0104*, *spd_1874* and *trpD* decreased by 2.7, 2.4, and 2.8 fold, respectively, in the Δ*cvfD* mutant with respect to the WT strain (Fig. S3). The relative amounts of the *pstS*, *phoU1*, *psaA, czcD*, *spd_0104* and *spd_1874* transcripts were restored to near-WT levels and *trpD* mRNA abundance was partially restored in the *cvfD*^+^-complemented strain background (Fig. 3B, 3C, 4A, 4B, and S3). This result confirms that the changes in the relative steady-state levels of these transcripts was caused by lack of CvfD.

**Figure 4:**
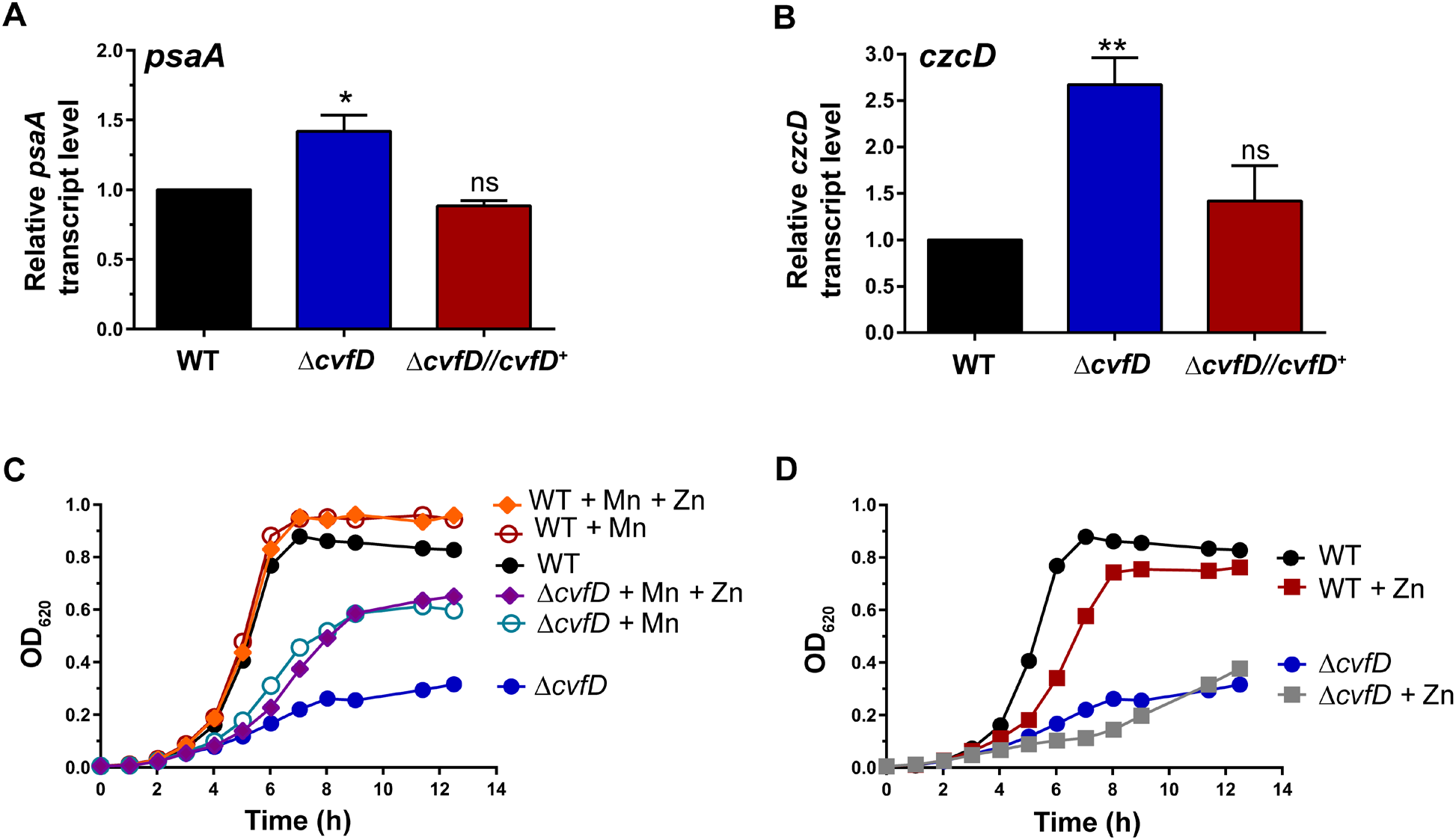
Loss of *cvfD* perturbs metal homeostasis in *S. pneumoniae* D39. (A, B) qRT-PCR analysis was used to determine *psaA* and *czcD* transcript levels in a wild-type D39 parent (WT; IU1781) and its derived isogenic mutants (Δ*cvfD*; IU4772 and Δ*cvfD bgaA*::P_*ftsA*_ *cvfD*^+^; IU5508) as described in the legend of Fig. 3. Data and error bars represent the means (± SEM) of at least three independent experiments and asterisks (*), (**), and ns indicate P<0.05, P<0.01, and not significantly different, respectively. (C) Growth characteristics of Δ*cvfD* (IU4772) mutant in BHI with no MnSO_4_ or ZnCl_2_ addition, addition of 0.5 mM MnSO_4_ (+ Mn), or addition of 0.5 mM MnSO_4_ and 0.2 mM ZnCl_2_ (+ Mn + Zn) relative to the wild-type parent (WT; IU1781) at 37°C. At least four independent growth curves were performed under the above-mentioned conditions except for the condition where both MnSO_4_ and ZnCl_2_ were added (+ Mn + Zn) for which two independent experiments were performed, and a representative curve is shown. (D) Growth characteristics of Δ*cvfD* (IU4772) mutant in BHI in the presence or absence of 0.2 mM ZnCl_2_ relative to the wild-type parent (WT; IU1781) at 37°C. A representative curve from at least six independent growth experiments is shown. (C, D) Average growth rates and growth yields are listed in Supplemental Table S3.

### CvfD mediates the abundance of small regulatory RNAs

Recent work in serotype 4 strain (TIGR4) shows that small regulatory RNAs (sRNAs) play crucial roles in *S. pneumoniae* pathogenesis (48). Based on these findings, we tested whether CvfD played any role in regulating relative sRNA transcript amounts in *S. pneumoniae* D39, where the presence of over one-hundred sRNAs have been reported by recent studies (49, 51, 65). sRNA transcript levels in replicates of a second Δ*cvfD* mutant construct (NRD10133; Table S1 and S2) were compared to that in its WT parent strain (IU1781) using small RNA-seq (sRNA-seq) analysis of total RNA extracted from exponentially growing cultures at OD_600_ ≈0.15 in matched batches of BHI broth (see *Materials and Methods*). sRNA-seq analysis revealed twenty sRNAs that showed a 1.8 fold or greater difference in relative amount between the Δ*cvfD* mutant and WT strain (Table 2). Interestingly, seven of the sRNAs that were down-regulated in the Δ*cvfD* mutant, included 3 Ccn sRNAs (CcnA, CcnD and CcnE) and Spd-sr85, which is a 6S RNA homolog (66–68). CcnA, CcnB, CcnC, CcnD, and CcnE are a set of homologous, highly conserved pneumococcal sRNAs that are under positive transcriptional control of the CiaR response regulator and function to inhibit competence development via base-pairing with the precursor of the competence stimulatory peptide-encoding mRNA, *comC* (49, 69, 70). mRNA-seq analysis reveals that relative *ciaR* transcript level is down-regulated by ≈2 fold in a Δ*cvfD* mutant (Table1), which could explain the subsequent down-regulation of the Ccn sRNAs. New results as part of a Grad-seq analysis of the TIGR4 strain indicate that its CvfD homolog (SP_1537) may also bind weakly to CcnB, but not to other Ccn sRNAs (53). Thirteen sRNA transcripts were up-regulated in the Δ*cvfD* mutant, including two anti-toxin sRNAs (Spd_sr110 and Spd_sr112), the previously characterized sRNA Spd_sr37, and a toxin RNA encoded by Spd_sr23 (Table 2). Together, these findings show that CvfD mediates sRNA amounts in *S. pneumoniae*, possibly by regulating sRNA gene expression.

**Table 2:** Relative sRNA transcript level changes in strain a Δ*cvfD* mutant compared to the *cvfD*^+^ parent strain under exponential growth in BHI broth^a^.

| sRNA ID | Flanking genes | Fold change | P<sub>adj</sub> value |
| --- | --- | --- | --- |
| Decreased relative expression |  |  |  |
| CcnE | *spd_0221, spd_0222* | -2.38 | 2.60E-07 |
| CcnA | *spd_0240, ruvB* | -2.38 | 0.006544 |
| CcnD | *spd_0242, uppS* | -1.89 | 1.69E-05 |
| Spd_sr50 | *spd_0867, spd_0868* | -3.85 | 1.41E-21 |
| Spd_sr65 | *spd_1161, spd_1160* | -2.22 | 0.001549 |
| Spd_sr82 | *spd_1455, spd_1454* | -2.04 | 8.18E-08 |
| Spd_sr83 | *recG, spd_1506* | -7.14 | 2.61E-27 |
| Increased relative expression |  |  |  |
| Spd_sr13 | *spd_0101, capD* | 2.41 | 1.94E-15 |
| Spd_sr18 | *spd_0130, gidA* | 2.01 | 6.10E-13 |
| Spd_sr23 | *spd_0240, ruvB* | 2.93 | 3.97E-25 |
| Spd_sr37 | *tmrU, spd_0128* | 2.04 | 2.31E-12 |
| Spd_sr61 | *spd_1080, spd_1079* | 3.63 | 4.70E-38 |
| Spd_sr66 | *spd_1175, spd_1174* | 4.72 | 0.003515 |
| Spd_sr67 | *spd_1180, spd_1179* | 3.12 | 3.22E-13 |
| Spd_sr73 | *spd_1289, spd_1288* | 2.75 | 1.57E-15 |
| Spd_sr90 | *spd_1664, treC* | 2.53 | 2.96E-14 |
| Spd_sr101 | *spd_1834, spd_1833* | 5.13 | 1.43E-27 |
| Spd_sr107 | *malP, spd_1931* | 3.66 | 2.61E-27 |
| Spd_sr110 | *adcA, spd_1996* | 4.59 | 3.14E-37 |
| Spd_sr112 | *adcA, spd_1996* | 5.66 | 3.75E-54 |

<sup>a</sup>RNA extraction and sRNA-seq analyses were performed as described in *Materials and Methods*. RNA was prepared from cultures of the encapsulated parent strain IU1781 (wild-type parent; D39 *rpsL1 cvfD*<sup>+</sup>) and its derived mutant NRD10133 (D39 *rpsL1 ΔcvfD*) (Table S1 and S2). Fold changes (1.8-fold cut-off) and P-values ( $P_{\text{adj}}$ value <0.05) are based on three independent biological replicates.

### CvfD functions as a post-transcriptional regulator of the dual phosphate transport system in *S. pneumoniae*

The list of genes whose transcript levels were significantly altered in the Δ*cvfD* mutant relative to the WT strain revealed that many are members of multi-gene operons (Table 1). Of special note is the *pst1* (phosphate transport system 1) operon that is maximally upregulated by ≈4 fold in the Δ*cvfD* mutant compared to the WT strain. *pst1* is one of two distinct phosphate ion (Pi) transporters in *S. pneumoniae* D39, but it is not normally expressed in the WT strain during growth in BHI broth (71). In BHI broth, which contains a relatively high concentration of Pi, the *pst2* operon, which encodes a second Pi transporter, is constitutively expressed, including the PhoU2 master regulator. PhoU2 negatively regulates *pst1* operon expression by blocking transcription activation by the phosphorylated PnpR~P response regulator (71). Consequently, *pst1* expression is derepressed in a *phoU2* deletion mutant (71). We observed that the Δ*cvfD* mutant phenocopies a Δ*phoU2* mutant at the transcriptional level, in that the same set of transcripts change relative expression in the Δ*cvfD* mutant as in the Δ*phoU2* mutant, albeit to different folds (Table 3). Yet, the transcript steady-state levels of *phoU2* and *pst2* polycistronic mRNA are not reduced in the Δ*cvfD* mutant compared to the wild-type (WT) strain (Table 1 and Fig. 3D). Therefore, we hypothesized that the absence of CvfD post-transcriptionally reduces PhoU2 amount, resulting in increased *pst1* expression.

**Table 3:**
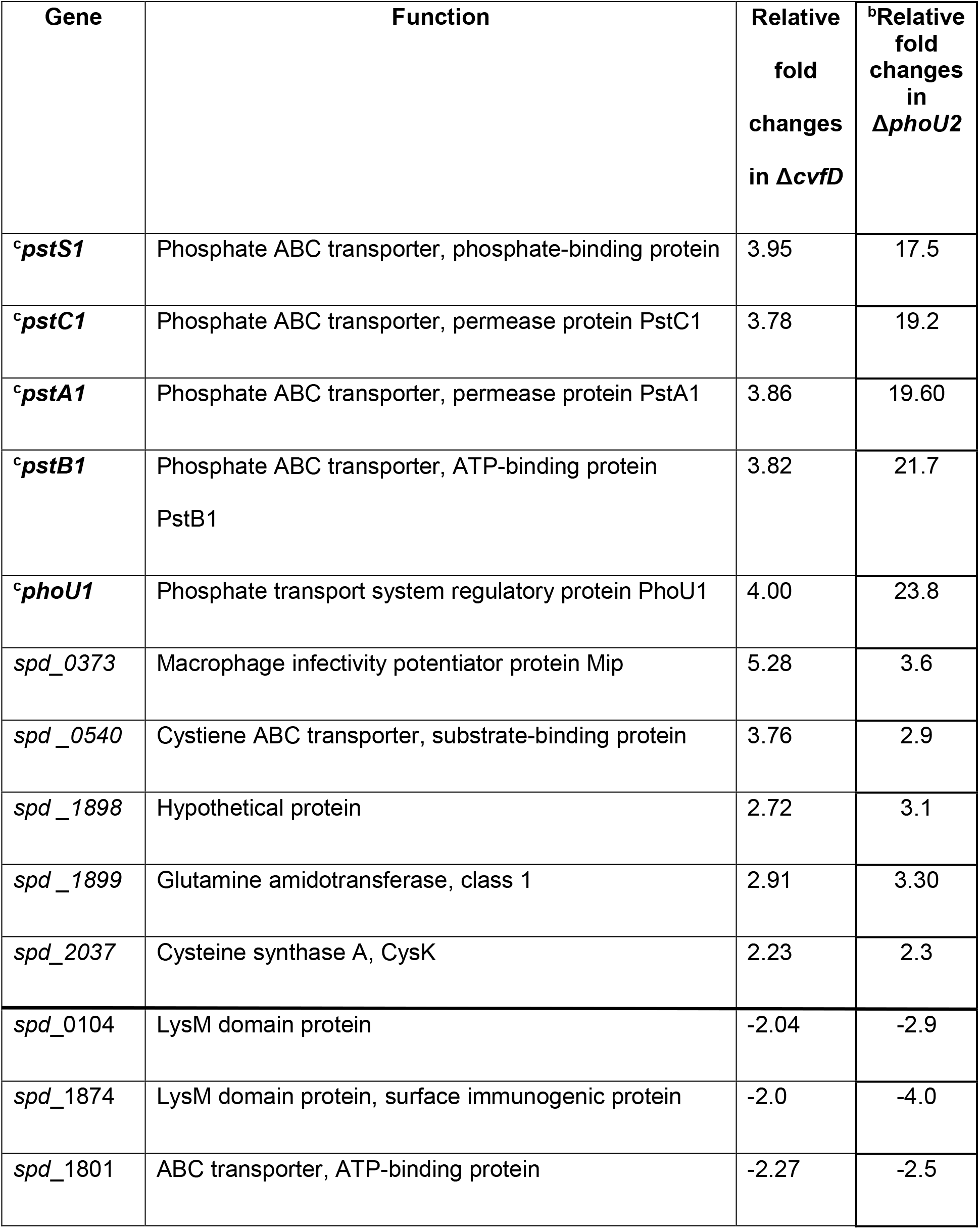
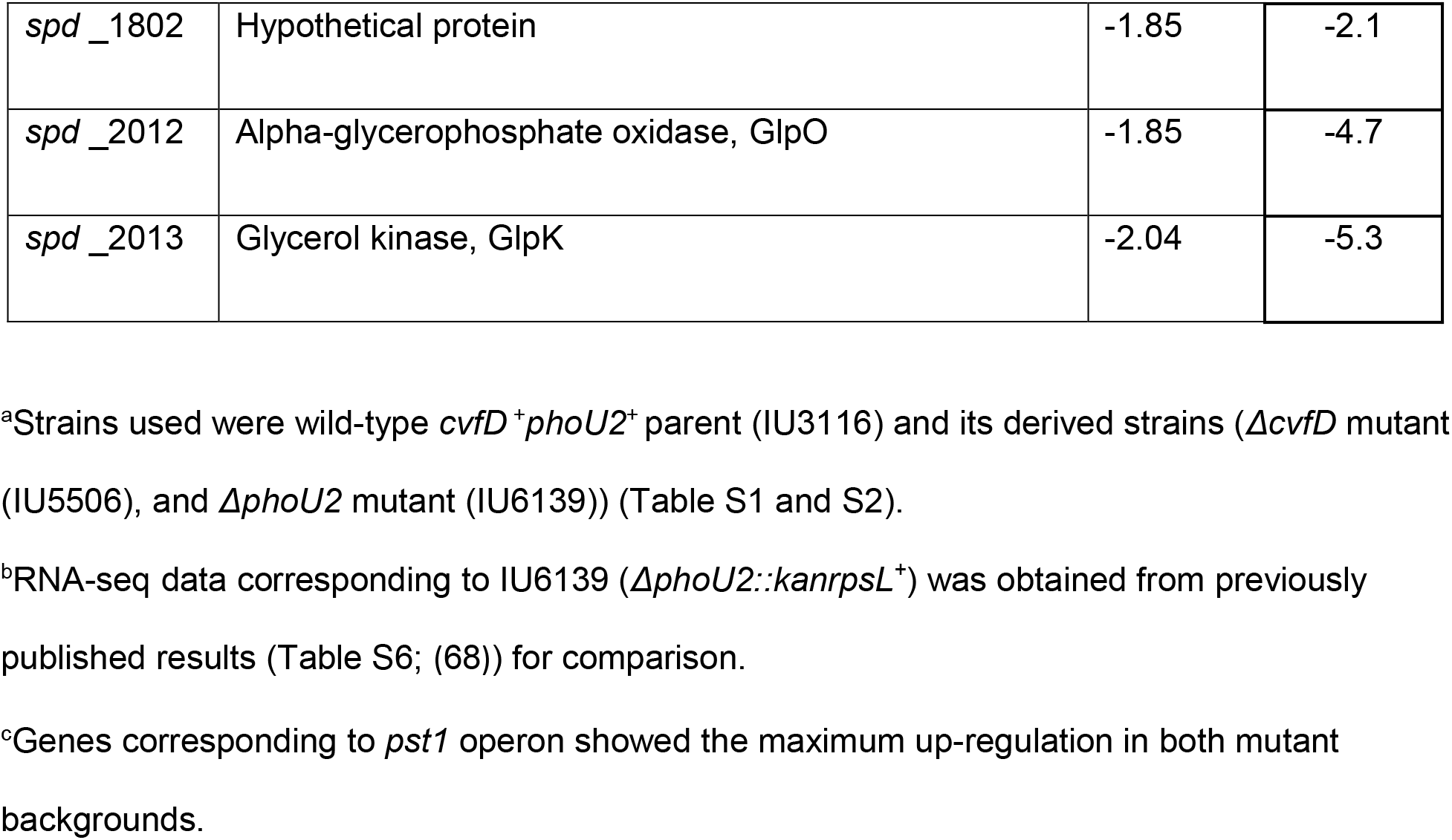
Relative transcript changes common between Δ*cvfD* and Δ*phoU2* mutants compared to the parent (*cvfD*^+^*phoU2*^+^)^a^.

| **Gene** | **Function** | **Relative fold changes in $\Delta cvfD$** | **<sup>b</sup>Relative fold changes in $\Delta phoU2$** |
| --- | --- | --- | --- |
| <sup>c</sup>*pstS1* | Phosphate ABC transporter, phosphate-binding protein | 3.95 | 17.5 |
| <sup>c</sup>*pstC1* | Phosphate ABC transporter, permease protein PstC1 | 3.78 | 19.2 |
| <sup>c</sup>*pstA1* | Phosphate ABC transporter, permease protein PstA1 | 3.86 | 19.60 |
| <sup>c</sup>*pstB1* | Phosphate ABC transporter, ATP-binding protein PstB1 | 3.82 | 21.7 |
| <sup>c</sup>*phoU1* | Phosphate transport system regulatory protein PhoU1 | 4.00 | 23.8 |
| *spd_0373* | Macrophage infectivity potentiator protein Mip | 5.28 | 3.6 |
| *spd_0540* | Cystiene ABC transporter, substrate-binding protein | 3.76 | 2.9 |
| *spd_1898* | Hypothetical protein | 2.72 | 3.1 |
| *spd_1899* | Glutamine amidotransferase, class 1 | 2.91 | 3.30 |
| *spd_2037* | Cysteine synthase A, CysK | 2.23 | 2.3 |
| *spd_0104* | LysM domain protein | -2.04 | -2.9 |
| *spd_1874* | LysM domain protein, surface immunogenic protein | -2.0 | -4.0 |
| *spd_1801* | ABC transporter, ATP-binding protein | -2.27 | -2.5 |
| *spd*_1802 | Hypothetical protein | -1.85 | -2.1 |
| *spd*_2012 | Alpha-glycerophosphate oxidase, GlpO | -1.85 | -4.7 |
| *spd*_2013 | Glycerol kinase, GlpK | -2.04 | -5.3 |

<sup>a</sup>Strains used were wild-type *cvfD*<sup>+</sup>*phoU2*<sup>+</sup> parent (IU3116) and its derived strains ( $\Delta$ *cvfD* mutant (IU5506), and $\Delta$ *phoU2* mutant (IU6139)) (Table S1 and S2).
<sup>b</sup>RNA-seq data corresponding to IU6139 ( $\Delta$ *phoU2::kanrpsL*<sup>+</sup>) was obtained from previously published results (Table S6; (68)) for comparison.
<sup>c</sup>Genes corresponding to *pst1* operon showed the maximum up-regulation in both mutant backgrounds.

To test this hypothesis, we fused an HA epitope tag to the C-terminal end of PhoU2 expressed at its chromosomal locus. The strains expressing PhoU2-HA did not exhibit growth defects caused by a Δ*phoU2* mutant (71), indicating that PhoU2-HA is functional (Fig. 5A; Table S3). Next, we introduced Δ*cvfD* into the strain expressing PhoU2-HA. PhoU2 expression was determined in the Δ*cvfD* mutant (Δ*cvfD* PhoU2-HA) and the parent (*cvfD*^+^ PhoU2-HA) strain by western blotting analysis. The Δ*cvfD* mutant contains 3-4 fold less PhoU2-HA than the WT parent, and PhoU-HA amount is restored to nearly WT levels when CvfD is expressed ectopically from a constitutive promoter in the Δ*cvfD* PhoU2-HA strain (Fig. 3E and 3F).

**Figure 5:**
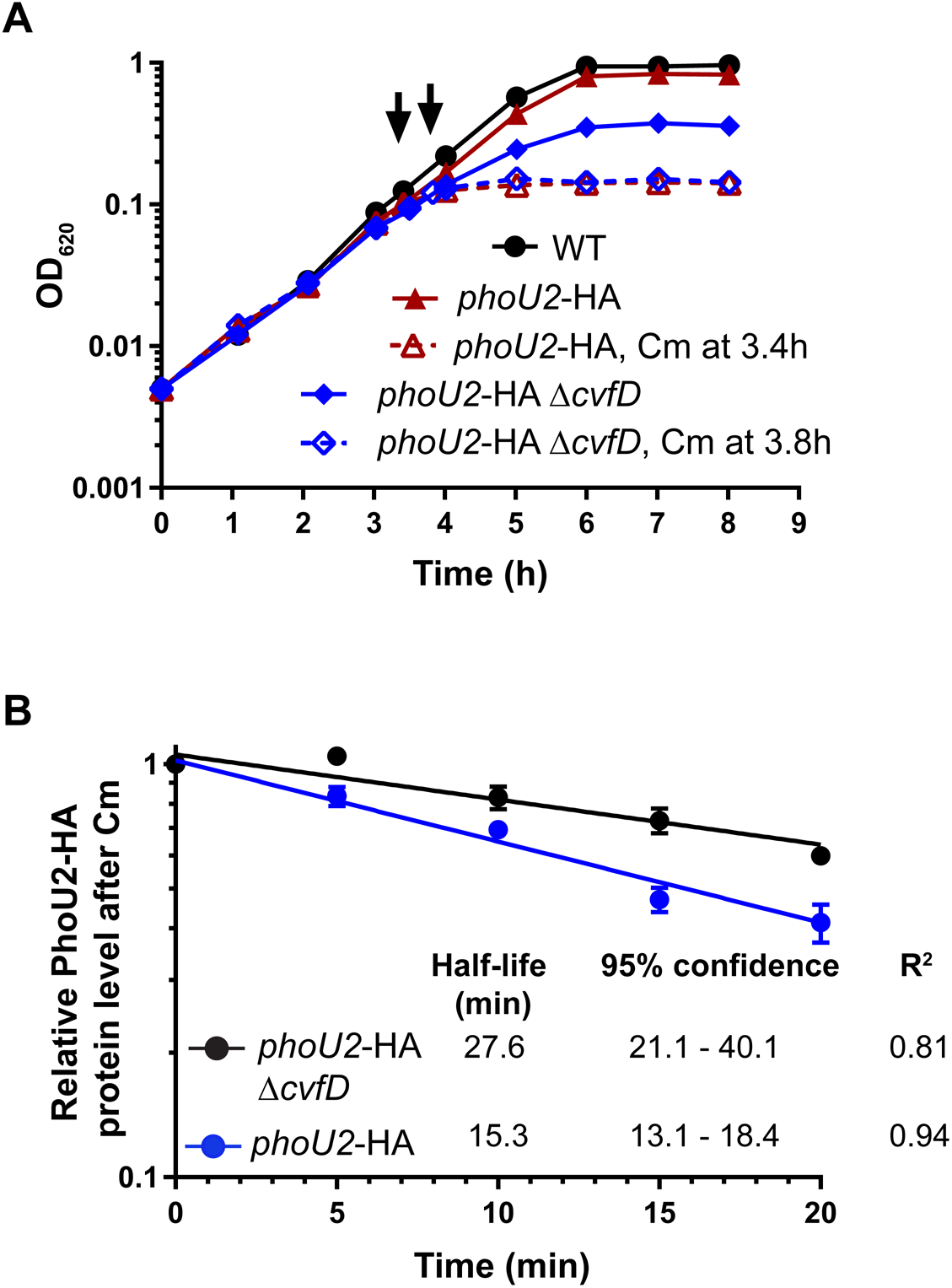
PhoU2-HA protein is more stable in a Δ*cvfD* strain than in a *cvfD*^+^ strain. (A) Representative growth and chloramphenicol (Cm) response curves of encapsulated D39 parent strain (IU1781, WT), IU8675 (*phoU2*-HA) and IU8722 (*phoU2*-HA Δ*cvfD*) strains. Arrows indicate addition of Cm at OD_620_ ~0.1. (B) Semi-log decay plot of relative PhoU2-HA protein amounts vs time after Cm addition obtained from three independent experiments. Half-lives of PhoU2-HA proteins in IU8675 (*phoU2*-HA) and IU8722 (*phoU2*-HA Δ*cvfD*) were determined after treatment with chloramphenicol (Cm), a protein translation inhibitor, as described in Material and Methods. Each data point represents the mean ± SEM (where not visible, error bars are smaller than the symbol) of PhoU2-HA protein at each time point after Cm treatment. Half-life values, 95% confidence intervals of half-life values and R^2^ value of the decay curves were obtained from non-linear regression analysis with GraphPad Prism as described in *Materials and Methods*. The amount of PhoU2-HA protein in *phoU2*-HA Δ*cvfD* at T_0_ relative to that in *phoU2*-HA *cvfD*^+^ strain at T0 was determined to be 0.48 ± 0.03 (mean ± SEM) from three independent experiments (See Fig. S4).

Lack of change in *phoU2* transcript amount and reduction of PhoU2 protein amount (Fig. 3D and 3F) in a Δ*cvfD* mutant suggests that either CvfD is a positive regulator of PhoU2 translation or a negative regulator of PhoU2 proteolysis. To distinguish between these two explanations, we determined the half-life of PhoU2-HA in the WT and Δ*cvfD* mutant following chloramphenicol addition to block translation (Fig. 5 and S4). To take rapid time points following chloramphenicol addition, a different protein extraction method was used for time-course quantitative western blotting (Fig. S4) than for western blots of steady-state cultures (Fig. 3E and 3F). In this different procedure, relative cellular PhoU2-HA amount was consistently 2 fold lower in the Δ*cvfD* mutant than the parent strain (Fig S4C, lanes 3 and 8). However, the half-life of PhoU2-HA was increased by ≈1.8 fold, rather than decreased, in the Δ*cvfD* mutant compared to the WT strain (Fig. 5B). Thus, we conclude that CvfD protein acts as a post-transcriptional positive regulator of PhoU2 translation.

Last, we used co-immunoprecipitation (co-IP) experiments to determine if CvfD interacts with the *phoU2* transcript. A strain expressing CvfD-FLAG^3^ was grown exponentially in BHI broth, and CvfD-FLAG^3^ was collected from cell lysates by binding to anti-FLAG magnetic beads (see *Materials and Methods*). RNA was isolated from CvfD-FLAG^3^ eluted from the beads and subjected to qRT-PCR to detect co-immunoprecipitated *phoU2* transcript, where an extract for the parent strain expressing WT untagged CvfD was used as the control (Fig. 3G). *phoU2* transcript amounts were appreciably enriched by ≈4 fold from the strain expressing CvfD-L-FLAG^3^ compared to WT strain. No significant difference was observed in the amount of 16S rRNA (control RNA) that co-immunoprecipitated in strains expressing WT CvfD^+^ or CvfD-L-FLAG^3^ (Fig. 3G). We conclude that CvfD acts as a positive regulator of PhoU2 translation by binding to *phoU2* mRNA.

### Null mutations in capsule synthesis suppress the requirement for CvfD in *S. pneumoniae* D39

Δ*cvfD* mutants occasionally acquire spontaneous mutations that restore growth rate and yields in aged BHI broth or at 32 °C (Fig. 2B and S5; Table S3 and S4). Two independently isolated spontaneous Δ*cvfD* growth suppressors (IU7291 and IU7293; Table S1, *Materials and Methods*) and one independently isolated Zn(II)-stress growth suppressor (IU7294; Table S1; see below) were stored for further characterization. Each suppressed Δ*cvfD* mutant shows WT growth rate and increased growth yield at 37 °C and 32°C compared to its Δ*cvfD* parent (Fig. S5; Table S3). Whole-genome sequencing showed that Δ*cvfD* suppressor isolates 1 and 3 contain complex combinations of mutations (Table S4). Isolate 1 contains a deletion that extends into the *spxB* gene encoding pyruvate oxidase that produces acetyl-phosphate and hydrogen peroxide (72–74); however, a constructed Δ*spxB* Δ*cvfD* double mutant (IU17114) shows the same growth defects in aged BHI broth as its Δ*cvfD* parent (data not shown), indicating that Δ*spxB* is not the suppressor mutation. We did not deconvolute and test the other mutations in isolates 1 and 3 further in this study.

In contrast, isolate 2 contained a point mutation in *cps2E*, leading to a R307S change that likely abolishes the activity of the glycosyltransferase required for the initial step of pneumococcal type 2 capsule synthesis (75). To test this idea, a frameshift truncation mutation ΔA in codon 326 of *cps2E* (*cps2E*(ΔA)) was introduced in a Δ*cvfD* mutant background and the growth characteristic of the Δ*cvfD cps2E*(ΔA) double mutant was determined at 37°C and 32°C. The growth characteristics of the Δ*cvfD cps2E*(ΔA) double deletion mutant are comparable to those observed for the WT *cvfD*^+^ *cps*^+^ and *cvfD^+^ cps2E*(ΔA) strains, confirming that lack of capsule is sufficient to suppress the growth defects of a Δ*cvfD* mutant (Fig. 6; Table S3). Notably, the *cps2E*(ΔA) mutant eliminated the strong CS phenotype of the Δ*cvfD* mutation on TSAII BA plates (Fig. S2B). The accumulation of capsule synthesis mutations that suppress Δ*cvfD* mutations may partly reflect the inverse relationship between phosphate transport and the capacity to synthesize capsule reported previously (see *Discussion*; (71)).

**Figure 6:**
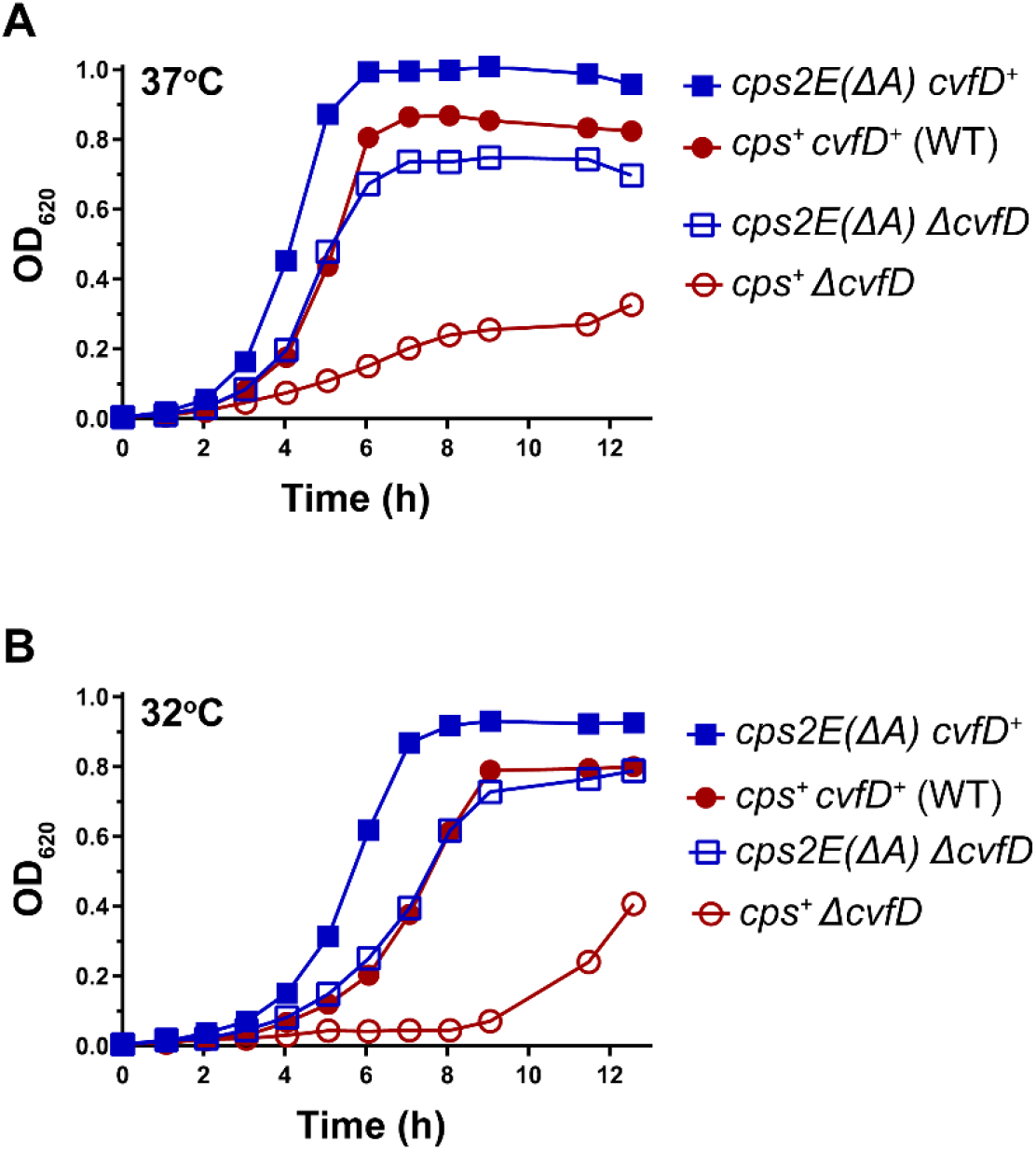
Loss of capsule suppresses the observed growth defect of a Δ*cvfD* mutant. Growth characteristics of the unencapsulated D39 *cps2E*(ΔA) strain (IU3309), encapsulated D39 *cps*^+^ *cvfD*^+^ parent strain (IU1781), and their derived mutant strains Δ*cvfD cps2E*(ΔA) (IU8396) and *cps*^+^ Δ*cvfD* (IU4772) at 37°C (A) and at 32°C (B). (A, B) At least two independent growth curves were performed at 37°C or 32°C, and a representative curve is shown. Average growth rates and growth yields are listed in Supplemental Table S3.

### CvfD regulates zinc and manganese homeostasis

We observed via RNA-seq analysis that the Δ*cvfD* mutant strain showed an up-regulation of *psaBCA* operon encoding the manganese (Mn (II)) uptake system and *czcD* encoding the zinc (Zn (II)) efflux pump by ~2 fold and ~4 fold respectively (Table 1). Furthermore as shown in Figures 4A and 4B, CvfD dependent up-regulation of *psaA* and *czcD* was confirmed by qRT-PCR analysis. Previous work by Jacobsen et al. (2011) has shown that Zn (II) toxicity as indicated by the up-regulation of *czcD* results in Mn (II) deficiency leading to subsequent up-regulation of the *psaBCA* operon (76). Moreover, relative transcript levels of *nmlR,* the gene encoding nitric oxide (NO) detoxification system, which increases in expression upon Zn stress (76), was also up-regulated by ≈3 fold in the Δ*cvfD* mutant compared to a WT strain further indicating that deletion of *cvfD* results in cytoplasmic Zn stress in *S. pneumoniae*. Consistent with this idea, addition of 500 μM Mn^2+^ to aged BHI substantially restored the growth of a Δ*cvfD* mutant (Fig. 4C; Table S3). In contrast, addition of excess 200 μM Zn^2+^ reduced the Δ*cvfD* mutant yield phenotype compared to cultures lacking additional Zn^2+^ (Fig. 4D, Table S3) and occasionally led to the accumulation of faster growing suppressors (Tables S1 and S4; Fig. 4D). However, addition of both 500 μM Mn^+2^ and 200 μM Zn^2+^ increased the growth yield and restored the growth rate of the Δ*cvfD* mutant to that of the WT strain (Fig. 4C; Table S3). Finally, we recapitulated the experiments in aged BHI in a chemically defined medium lacking added Mn^2+^ (CDM-Mn) (*Materials and Methods*). There is sufficient Mn^2+^ trace contamination in CDM-Mn to allow nearly normal growth of the WT strain (Fig. S6; Table S3). Similar to aged BHI, the Δ*cvfD* mutant grew much slower and to a lower yield than the WT strain in CDM-Mn, and the addition of 500 μM Mn^2+^ restored the growth rate and yield to the WT level, even when 200 μM Zn^2+^ was added (Fig. S6; Table S3). These results suggest that aged BHI and CDM-Mn media lack sufficient available Mn^2+^ ion concentration to support growth of Δ*cvfD* mutants and to prevent likely Zn^2+^ toxicity.

## DISCUSSION

RNA-binding proteins regulate gene expression at the post-transcriptional level by variety of mechanisms. In this study, we characterized the S1 RNA-binding domain protein CvfD that is highly conserved among the *Firmicutes* (Fig. 1). We identified this RNA-binding protein as a pleiotropic regulator in *S. pneumoniae* that controls the relative transcript levels of at least 144 mRNAs and sRNAs involved in diverse cellular processes (Tables 1 and 2). Furthermore, we show that besides causing global changes in gene expression, deletion of *cvfD* results in several phenotypes, including reduced growth rate and yield in some media (Fig. 2A, 4C, 4D, and S6; Table S3), cold-sensitivity (Fig. 2B and S2), reduced virulence (Fig. 2F), manganese deficiency (Fig. 4C and S6), and likely zinc toxicity (Fig. 4D). Our investigation into the role of CvfD in the regulation of phosphate transport system 1 (Pst1) revealed that this protein is a post-transcriptional regulator that binds and promotes the translation of the transcript encoding PhoU2 (Fig. 3), a master regulator of phosphate transport.

Among the 144 genes that display altered transcript expression in the Δ*cvfD* mutant compared to the WT strain are 124 that encode proteins and 20 that synthesize sRNAs (Fig. 3A; Tables 1 and 2). Notably, among the transcript amounts down-regulated in the Δ*cvfD* mutant are the *piu* operon encoding the iron uptake system (*piuC*, *piuD*, *piuA*), the *trp* operon encoding genes involved in tryptophan biosynthesis (*trpABFCDGE*, *trpY*), and the *ciaRH* operon encoding a TCS involved in biofilm development (77), competence (78), and virulence (79). Consistent with decreased expression of CiaRH, transcripts positively regulated by this TCS (*htrA-parB*, *axe*, *spd_0537*, *ccnA, ccnD, and ccnE*) are down-regulated (Table 1 and 2). Relative transcript amounts of genes encoding extracellular LysM proteins (*spd_0104* and *spd_*1874), which are positively regulated by the cell-wall stress TCS WalRK, are also downregulated in the Δ*cvfD* mutant, as is *dltA*, which D-alanylates lipoteichoic acid and imparts resistance to antimicrobial peptides (80, 81). Among the up-regulated transcripts in the *cvfD* deletion strain are mRNAs encoding proteins involved in phosphate transport (*pstS1*, *pstC1*, *pstA1*, *pstB1*) (Fig. 3B and 3C), manganese uptake (*psaB*, *psaC*, *psaA*), zinc efflux (*czcD*), vitamin B1 metabolism (*thiW*, *thiM-2*, *thiE-2*), vitamin B6 biosynthesis (*pdxT*, *pdxS*), sugar catabolism (*lacD*, *lacC*, *lacB*, *lacA*, *scrB*, *nagA*, *pgk*, *gap*), general stress responses, (*hrcA*, *grpE*, *nrdD*, *groEL*, *phtA*), and virulence (*mip, prtA*, *pspA*, *phtA*) (Table 1). We conclude that lack of CvfD is highly pleiotropic impacting the transcripts of numerous key metabolic and virulence operons of *S. pneumoniae* (Fig. 7).

**Figure 7:**
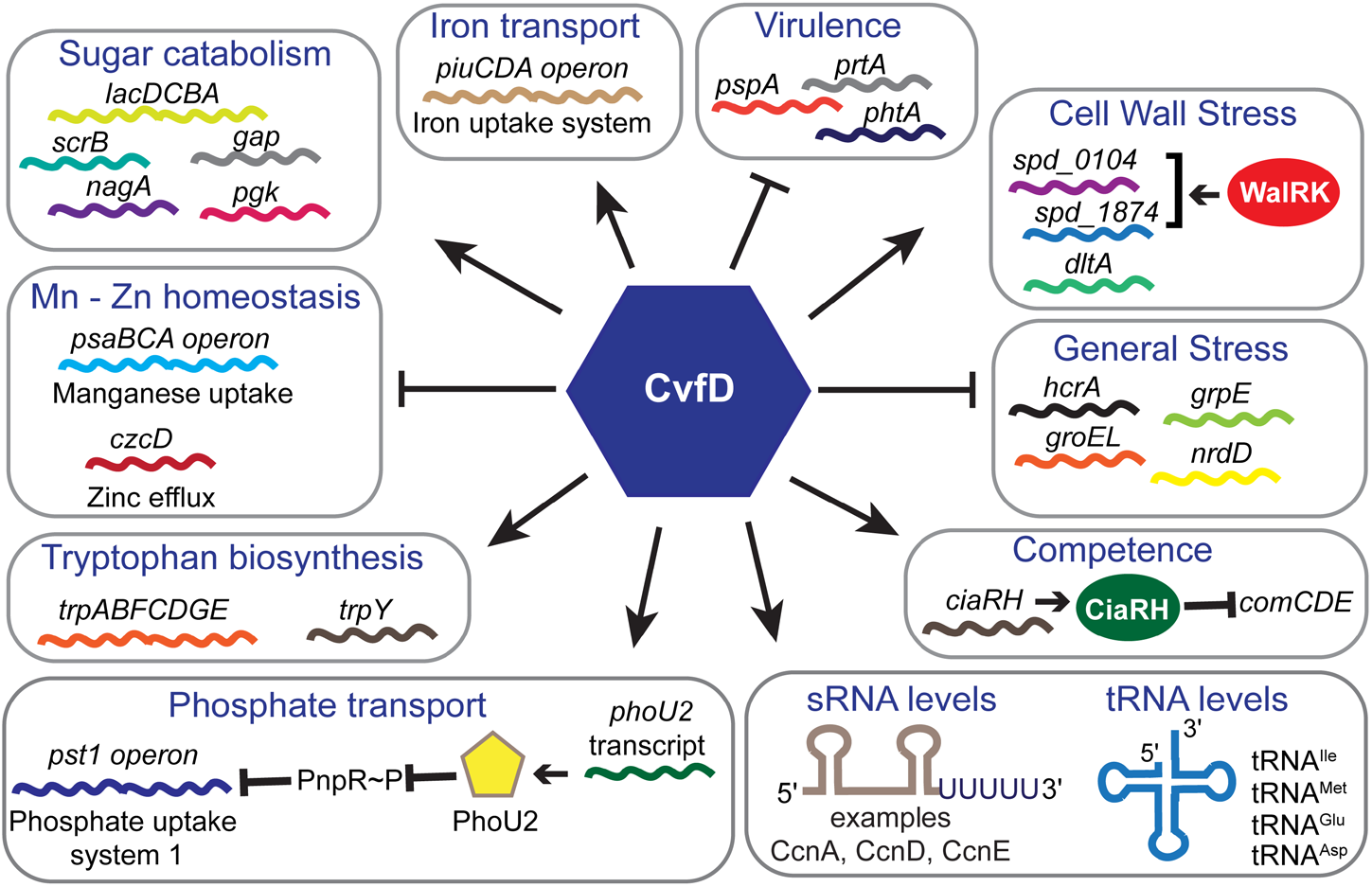
Overview of the regulatory scope of CvfD in *S. pneumoniae*. Major pathways that are regulated by CvfD are indicated.

The mechanisms underlying these effects of CvfD on transcript amounts remain to be determined. Some of the effects reflect downstream responses and may indirect. For example, Δ*cvfD* mutants showed growth rate and yield defects relative to the WT strain in aged BHI (Fig. 2A and S1) and in CDM-Mn media (Fig. S6), but not on TSAII BA plates (Fig. S2A). These growth defects are reversed by the addition of Mn^2+^ ion to the media. The deficiency of Mn^2+^ in *cvfD* mutants is reflected by the increased transcription of the *psaB, psaC,* and *psaA* manganese uptake ABC transporter and the increase of the *czcD* zinc efflux pump to ward off zinc toxicity by mismetallation (76, 82). However, it remains to be determined whether CvfD positively regulates translation of the PsaABC transporter proteins directly, such that the absence of CvfD results in a manganese deficiency, or whether the manganese deficiency is caused by a more indirect mechanism. Moreover, a recent Grad-seq study identified the CvfD homolog (SP_1537) in the TIGR4 strain as an RNA-binding protein that sediments in the region of the glycerol gradient and pellet indicative of general mRNA binding (53), which is reflected by the broad pleiotropy of Δ*cvfD* mutants.

Likewise, the absence of CvfD could decrease virulence of *S. pneumoniae* in several ways. Many of the genes whose transcription is altered in Δ*cvfD* mutants impact pneumococcal virulence. Some encode established pneumococcal virulence factors (e.g., *mip, prtA*, *pspA*, *phtA*, *dltA*), many mediate ion transport (e.g., iron, manganese, and phosphate) important in virulence (43, 83, 84), and others (e.g., the regulons of the CiaRH and WalRK TCSs) likely reflect changed peptidoglycan composition (52, 85). The cold-sensitivity in the absence of CvfD (Fig. 2B and S2) may also attenuate virulence, since the temperature of the nasopharynx (≈34 °C) colonized by *S. pneumoniae* is several degrees lower than that of the body core (≈37 °C) (62). Moreover, cellular CvfD amount is up-regulated moderately at the lower temperature of 32 °C compared to 37 °C, likely by a post-transcriptional mechanism (Fig. 2C-2E). Additional experiments are needed to understand how CvfD contributes to cold tolerance and virulence.

In this paper, we studied one case of CvfD regulation in detail. We show that CvfD is an RNA-binding protein, which post-transcriptionally regulates translation of PhoU2, a master regulator of phosphate transport in *S. pneumoniae* (71). Data herein shows that that CvfD interacts with the *phoU2* mRNA and positively regulates PhoU2 expression by increasing the translation of *phoU2* mRNA (Fig. 3, 5, and S4). Interestingly, Δ*cvfD* mutants readily accumulated mutations defective in capsule biosynthesis leading to faster growth (Fig. 6, S2, and S5; Table S3 and S4), and the selection for capsule loss may partly reflect a relationship between phosphate transport and capsule synthesis reported earlier (71). Encapsulated D39 requires a functional Pst1 or Pst2 transporter, making a Δ*pst1* Δ*pst2* double mutant synthetically lethal, which can be suppressed by loss of capsule synthesis (71). The accumulation of mutations in capsule synthesis that suppress growth defects of Δ*cvfD* mutants is consistent with the notion that down-regulation of PhoU2 in the absence of CvfD leads to intracellular phosphate stress that in turn interferes with capsule biosynthesis.

There are several mechanisms by which CvfD may be acting to positively regulate the expression of PhoU2. CvfD may facilitate *phoU2* translation by binding to the *phoU2* transcript and altering its RNA secondary structure. There are several known examples of RNA-binding proteins mediating translation activation. For example in *E. coli*, the RNA-binding protein CsrA activates translation of the *moaABCDE* operon encoding molybdenum biosynthesis genes by altering the conformation of the *moaA* transcript, without affecting its mRNA stability (86). Similarly, the CsrA ortholog RsmA of *P. aeruginosa* activates translation by binding and disrupting a secondary structure in target transcripts that would otherwise sequester ribosome binding sites (87). CvfD possesses a single conserved S1 RNA-binding domain, which is closely related to the cold-shock domains (CSD) found in cold-shock proteins that function to stabilize single-stranded regions of mRNAs, preventing the formation of unfavorable secondary structures that otherwise disrupt translation (88).

Alternatively, the S1 domain of CvfD could recruit ribosomes to specific target mRNAs, which in turn would improve translation efficiency and subsequently increase translation. This mechanism is analogous to the function of ribosomal protein S1, which works in complex with the 30S subunit of ribosomes to facilitate recognition of mRNAs for translation (89). Last, CvfD might mediate interactions between the *phoU2* transcript and an sRNA and/or another protein to activate *phoU2* translation, by analogy to the Hfq RNA chaperone in Gram-negative bacteria (21, 28). Future determinations of the CvfD binding and structural motifs in the *phoU2* and other target transcripts will distinguish among these mechanisms and serve as a model for how this S1-protein activates translation and possibly carries out other functions.

## MATERIALS AND METHODS

### Bacterial strains and growth conditions

Bacterial strains used in this study were derived from encapsulated *S. pneumoniae* serotype 2 strain D39W and are listed in Table S1. Strains were grown on plates containing trypticase soy agar II (modified; Becton-Dickinson [BD]) and 5% (vol/vol) defribrinated sheep blood (TSAII BA) at 37°C in an atmosphere of 5% CO_2_. Liquid cultures were grown statically in BD brain heart infusion (BHI) broth or a chemically defined medium (90) lacking added Mn^2+^ (CDM-Mn) at 37°C in an atmosphere of 5% CO_2_. Bacteria were inoculated into BHI broth from frozen cultures or single colonies. For overnight cultures, strains were first inoculated into a 17-mm-diameter polystyrene plastic tube containing 5 mL of BHI broth and then serially diluted by 100-fold into five tubes; these cultures were then grown for 10 to 16 h. Cultures with an optical density at 620 nm (OD_620_) of 0.1 to 0.4 were diluted to a starting OD_620_ between 0.002 and 0.005 in 5 mL of BHI broth in 16-mm glass tubes. Growth was monitored by measuring OD_620_ using a Spectronic 20 spectrophotometer. For antibiotic selections, TSAII BA plates and BHI cultures were supplemented with 250 μg kanamycin per mL, 150 μg streptomycin per mL, or 0.3 μg erythromycin per mL. For growth in CDM-Mn, overnight cultures were first grown in BHI to mid-exponential phase (OD_620_ 0.1 - 0.3), centrifuged at 3000 X *g* for 10 min at room temperature to collect cell pellets, and then suspended in the same volume of CDM-Mn. Dilutions were made in 3 mL of CDM-Mn to obtain an OD_620_ of 0.1 and subsequently diluted 100-fold in pre-warmed CDM-Mn to start cultures for growth experiments.

### Construction and verification of mutants

Mutant strains were constructed by transformation of competent *S. pneumoniae* strains with linear PCR amplicons as described previously (72). DNA amplicons containing antibiotic resistance markers were synthesized by overlapping fusion PCR. *S. pneumoniae* cells were induced to competence by the addition of synthetic competence stimulatory peptide. Markerless deletions and replacements of the target genes were constructed using the *kan*^R^-*rpsL*^+^ (Janus cassette) allele replacement method as described previously (91). In the first step, the Janus cassette was used to disrupt target genes in an *rpsL1* (Str^R^) strain background, and transformants were screened for kanamycin resistance and streptomycin sensitivity. In the second step, the Janus cassette was replaced by a PCR amplicon containing the desired mutation or replacement lacking antibiotic markers, and the resulting transformants were screened for streptomycin resistance and kanamycin sensitivity. Final transformants were isolated as single colonies three times on TSAII BA plates containing antibiotics listed in Table S1 and subsequently grown for storage in BHI containing the appropriate antibiotic. All constructs were confirmed by PCR amplification and sequencing.

Suppressor mutants IU7291, IU7292 and IU7294 were isolated as described below. After growth of strains IU4771 (Δ*cvfD*) and IU4708 (Δ*cvfD*::P_c_-[*kan*^R^-*rpsL*^+^]) in BHI broth for 12 h at 37°C, 400 μL aliquots were taken out, mixed with 600 μL of 25% glycerol and used to prepare temporary freezer stocks. The stored strains were first tested for their abilities to suppress the growth phenotype of a Δ*cvfD* mutant at 37°C. The temporary freezer stocks were then single colony isolated 3 times and stored as permanent frozen stocks IU7291, IU7293. IU7294 was similarly obtained from IU4772 (Δ*cvfD*) cultured in BHI broth containing 0.2 mM ZnSO_4_ for 12 h.

### RNA extraction

To isolate RNA for RNA-sequencing, strains were grown in 30 mL of BHI starting at an OD_620_ = 0.002 in 50 mL conical tubes. RNA was extracted from exponentially growing cultures of IU3116 (wild-type parent; D39 *rpsL1* CEP::P_c_-[*kan*^R^-*rpsL*^+^]) and its derived isogenic mutant IU5506 (D39 *rpsL1* Δ*cvfD CEP::* P_c_-[*kan*^R^-*rpsL*^+^]) at OD_620_ ≈ 0.1 for mRNA-seq analysis or IU1781 (wild-type parent; D39 *rpsL1*) and its derived markerless mutant NRD10133 (D39 *rpsL1* Δ*cvfD*) at OD_620_ ≈0.15 for sRNA-seq analysis using the FastRNA Pro Blue Kit (MP Bio) according to the manufacturer’s guidelines. Briefly, cells were collected by centrifugation at 14,500 x g for 5 min at 4°C. Cell pellets were resuspended in 1 mL of RNApro™ solution (MP Bio) and processed thrice in the Fast Prep Instrument (MP Bio) for 40 seconds at a speed setting of 6.0. Chloroform and 100% ethanol was added to extract and then precipitate RNA from the lysate, respectively. RNA purification was done using miRNeasy minikit (Qiagen), including on-column treatment with DNase I (Qiagen) following the manufacture’s guidelines. To isolate RNA for qRT-PCR, RNA was extracted from 5 mL of exponential phase cultures following the same method described above. 5 μg of total RNA was further digested with DNase using a DNA-free kit (Ambion). The amount and purity of all RNA samples isolated were assessed by NanoDrop spectroscopy (Thermo Fisher). RNA integrity of the samples used for RNA-seq library preparation was further assessed using the Agilent 2100 BioAnalyzer (Aligent Technologies).

### Library preparation and mRNA-seq

cDNA libraries were prepared from total RNA by the University of Wisconsin-Madison Biotechnology Center as described previously (51, 92). Briefly, total RNA was subjected to rRNA-depletion using RiboZero™ rRNA Removal Kit (EpiCentre Inc., Madison, WI, USA). Double stranded cDNA synthesis was performed with rRNA-depleted mRNA using ScriptSeq™ v2 RNA-Seq Library Preparation guide (EpiCentre Inc., Madison, WI, USA) in accordance with the manufacturer’s standard protocol. The amplified libraries were purified using Agencourt AMPure^®^ XP beads. Quality and quantity were assessed using an Agilent DNA 1000 chip (Agilent Technologies, Inc., Santa Clara, CA, USA) and Qubit® dsDNA HS assay kit (Invitrogen, Carlsbad, California, USA), respectively. Libraries were standardized to 2 μM and cluster generation was performed using standard Cluster kits (v3) and Illumina Cluster Station. Single-end 100 bp sequencing was performed using standard SBS chemistry (v3) on an Illumina HiSeq2000 sequencer. Images were analyzed using the standard Illumina pipeline, version 1.8.2.

### Library preparation and sRNA-seq

sRNA libraries were prepared from total RNA as described previously (51) with slight modifications. 5 μg of DNase-treated total RNA was first subjected to ribosomal RNA removal (RiboZero™ rRNA Removal for Gram-positive bacteria, Illumina). rRNA depleted samples were then subjected to RNA fragmentation using the Ambion RNA fragmentation kit (AM8740). Fragmented RNA was subjected to RNA 5’ polyphosphatase (Epicenter) treatment, which was performed to facilitate the 5’ adapter ligation step. Small RNA libraries were generated by Macrogen using TruSeq® small RNA library kit (Illumina). 100 bp paired-end read sequencing was performed using an Illumina HiSeq2500 sequencer.

### RNA-seq analysis

The raw sequencing reads were quality and adapter trimmed using Trimmomatic version 0.17 (93) with a minimum length of 90. The trimmed reads were mapped on the *Streptococcus pneumoniae* D39 (RefSeq NC_008533) genome and D39 plasmid pDP1 sequence (RefSeq NC_005022) using Bowtie2 (94). For mRNA-seq analysis custom PERL scripts were used to generate read counts for the genes and 100 bp non-overlapping intergenic regions of the genome. Differential gene expression was identified using the program DESeq2 (95) with these non-default parameters: estimateDispersions was set to “blind” and the sharingMode as “fit-only”. Genes were defined as differentially expressed if their P_adj_ (P-value adjusted for multiple testing) was less than 0.005.

Primary data from the mRNA-seq and sRNA-seq analyses were submitted to the NCBI Gene Expression Omnibus (GEO) and have the accession numbers GSE149546 and GSE148867, respectively.

### Quantitative RT-PCR (qRT-PCR) analysis

qRT-PCR analysis was performed as described previously (71, 92). 5 μg of purified RNA was treated by DNase from a DNA-free DNA removal kit (Ambion). 125 ng of treated RNA was used to synthesize cDNA by a qScript Felex cDNA synthesis kit (Quanta Biosciences). Synthesized cDNA was diluted 1:6 in water and then serially diluted 1:5 in water three more times. qRT-PCR reactions contained 10 μL of 2×Brilliant III Ultra-Fast SYBR Green QPCR Master Mix (Agilent), 2 μL of each 2 μM primers (Table S5), 0.3 μL of a 1:500 dilution of ROX reference dye, and 6 μL of diluted cDNA. Samples were run in an MX3000P thermocycler (Stratagene) with Program MxPro v. 3.0. Reactions were performed using cDNA from at least three independent biological samples and transcript amounts were normalized to 16S rRNA or *gyr*A RNA amount (internal control). Normalized transcript amounts were used to calculate fold changes of transcripts corresponding to target genes in different sets of mutants relative to the wild type parent. Statistical analysis was performed using one-way ANOVA (and nonparametric) in GraphPad Prism version 5.0.

### Mapping of suppressor mutations by whole genome sequencing

Strains IU7191, IU7193, and IU7194 containing mutations that suppressed the growth defect of a Δ*cvfD* deletion mutant were isolated as described in *Materials and Methods*. Overnight cultures were diluted into 5 mL of BHI broth to an OD_620_ of ~0.005 and grown to an OD_620_ of ~ 0.3 to 0.4. Cells were collected by centrifugation (10,000 X *g* for 10 min). Genomic DNA was purified from collected cells using a MasterPure Gram-positive DNA purification kit (Epicenter Biotechnologies) according to the manufacturer’s protocol. 1 μg of genomic DNA for each sample was diluted into 130 μL of TE and sheared using the following settings on the S220 focused-ultrasonicator (Covaris): 105W Peak Incident Power; 5 % Duty Factor, 200 Cycles per Burst; 40 seconds Treatment Time. Sheared samples were purified using AmpureXP beads (Agencourt/Seradyn/Beckman Coulter, A63881) to sample ratio of 0.5 X, and eluted with 54 μL of EB (Qiagen). Sheared samples were visualized using a D1K High Sensitivity tape (Agilent, 5067-5363) on an Agilent 2200 TapeStation, and consisted mostly of 650 to 700 bp fragments. Library construction was carried out on the Biomek FX^P^ (Beckman Coulter) using a modified SWHT (SPRIworks High Throughput for Illumina, Beckman Coulter) method to accommodate a 700 bp distribution. Bio Scientific NextFlex DNASeq Library Kit for Biomek FX^P^ (5140-42) was used in conjunction with Bioo Scientific NextFlex DNA barcodes-96 adaptors (514105). Following library construction, 15 μL of the 25 μL pre-enrichment library was used as template in a 10 cycle PCR amplification. Library sizes and concentrations were determined using TapeStation and Quant-iT Picogreen (Molecular Probes, P7589), respectively. Dilutions were made to 20 nM and pooled in preparation for a MiSeq run. 30 μL of the 20 nM library was diluted to 500 uL using Illumina buffer HT1 and loaded onto a 500 cycle MiSeq (version 2) flowcell (Illumina, MS-102-2003). Paired-end run cycle parameters were 260 (Read1) + 8 (Index read) + 260 (Read2). DNA sequences were assembled based on the published encapsulated D39 genome sequence (96). Bioinformatic analyses was performed using Cutadapt (https://code.google.com/p/cutadapt/) with a min length of 100 and quality cutoff of 30, and DNA sequence reads were assembled using newbler, mapped using bowtie, and called for SNPs using mpileup. Full coverage was obtained for the genomes of each mutant with >175 reads of most base pairs.

### Western Blotting

For data shown in Figures 2D and 3E, lysates were prepared from strains grown exponentially in BHI broth to OD_620_ of 0.1. Cells were collected by centrifugation (8,000 X *g* for 10 min at 4°C). Cell pellets were resuspended in 1 mL of cold 1 X PBS (4°C) containing protease inhibitor (ThermoFisher; cat# 78429) and were transferred into tubes containing lysing matrix B (MP Biomedicals, Inc.). Tubes were shaken in a FastPrep homogenizer (MP Biomedicals, Inc.) at 4°C for three 40s cycles at a setting of 6.0 M/s. Cell debris was removed by centrifugation at 16,000 X *g* for 5 min at 4°C. Approximately equal amounts of protein lysates normalized to OD_620_ were loaded per lane. Protein preparations for half-life studies were prepared using the SEDS method as described below. Samples were separated on 4 to 15% mini-protean TGX pre-cast SDS-PAGE gels (Bio-Rad, 456-1083) and subsequently transferred to nitrocellulose membranes. FLAG- and HA-tagged proteins were detected by Western blotting using a 1:1000 dilution of primary anti-FLAG rabbit polyclonal antibody (Sigma, F7425) or anti-HA rabbit polyclonal antibody (Invitrogen, 71-5500) at 1 μg mL^−1^, a 1:10000 dilution of HRP-conjugated donkey anti-rabbit antibody (GE Healthcare, NA934), and ECL detection reagent. Chemiluminescent signals were detected with an IVIS imaging system as described in (97).

### Half-life determination of PhoU2-HA

IU1781 (WT), IU8675 (*phoU2*-HA), and IU8722 (*phoU2*-HA Δ*cvfD*) strains were diluted from overnight cultures into 32 mL of fresh BHI in 50 mL conical tubes in the morning to OD_620_ = 0.005, and incubated at 37°C in an atmosphere of 5% CO_2_. A set of parallel 5-mL cultures in 13 mm-glass tubes were used to monitor the OD_620_ of the cultures before and after treatment with chloramphenicol (Cm) (Fig. 5A), which inhibits protein synthesis by preventing protein chain elongation. At OD_620_ ~0.1, 496 or 80 μL of 500 μg/mL Cm were added to 31 or 5 mL of IU8675 or IU8722 culture to obtain a final concentration of 8 μg/mL of Cm. 3 mL of culture from the 31-mL culture was removed immediately (t=0) into ice-prechilled 17-mm-diameter plastic tubes and swirled in ice for 30 sec, while the remaining culture was placed back into the incubator. Each chilled culture was placed into two prechilled microfuge tubes and centrifuged for 1.5 min at 16,000 X g at 4°C. The supernatants were removed, and the two pellets were resuspended in 1mL of PBS prechilled to 4°C and centrifuged for 1.5 min at 16,000 X g at 4°C. The supernatant was removed and the pellet was placed on dry ice. This procedure was repeated every 5 minutes to collect samples at 5, 10, 15 and 20 minutes after Cm treatment. The SEDS (0.1% deoxycholate, 150 mM NaCl, 0.2% SDS, 15 mM EDTA pH 8.0) lysis procedure was used to obtain protein for Western blot analysis, and total protein concentrations were determined (98). 6 μg of total protein of untreated IU1781 (non-HA tagged control) and strains IU8675 and IU8722 at various time after Cm treatment lysates were loaded on a 10% precast gradient SDS-PAGE gel (Bio-Rad), subjected to electrophoresis and transferred to a nitrocellulose membrane. PhoU2-HA proteins were detected with an anti-HA rabbit polyclonal antibody (Invitrogen, 71-5500, 1:500 dilution) as primary antibodies, and ECL anti-rabbit IgG horseradish peroxidase linked whole antibody as secondary (dilution 1:10,000). Chemiluminescent signals in protein bands were detected with an IVIS imaging system and quantitated as described in Wayne et al., 2010 (97) by subtracting the photon flux value of each signal band in a box with a background value of a box of the same size at the same blot location obtained with the non-HA tagged strain (IU1781) control. In order to ensure that chemiluminescent signals determined for the decay curves were within the linear range of detection with the Western procedure, 6, 4 and 2 μg of IU8675 lysate collected at T_0_ were loaded on each gel (lanes 2, 8 and 9 of Fig. S4A and lanes 8, 9 and 10 of Fig. S4C, respectively) and the signals from these lanes were used to generate a standard curve of chemiluminescent signal values vs protein amounts (Fig. S4B and Fig. S4D). The standard curve generated from each Western blot demonstrated that the chemiluminescent signals obtained with the Cm-treated samples (lanes 2 to 6) were in the linear range of Western detection for each experiment. The relative ratios of protein amounts after Cm treatment were extrapolated from the standard curve (shown as ratio under the bands on Fig. S4A and Fig. S4C). To obtain half-life values, normalized relative protein amounts at different times after Cm treatment from three independent experiments were plotted vs time on a semi-log curve, and half-life values of PhoU2-HA present in the *cvfD*^+^ or Δ*cvfD* strains, 95 % confidence values of the half-lives, and the R^2^ values of the non-linear regression curves were obtained from non-linear regression analysis with GraphPad Prism.

The ratios of PhoU2-HA present in the *cvfD*^+^ strain relative to the Δ*cvfD* strain at T_0_, or vice versa, were also determined with two western blot analyses (see lanes 2 and 10 of Fig. S4A, and lanes 3 and 8 of Fig. S4C) for each set of T_0_ samples. The amount of PhoU2-HA protein in *phoU2*-HA Δ*cvfD* at T_0_ relative to that in *phoU2*-HA *cvfD*^+^ strain at T_0_ was determined to be 0.48 ± 0.03 (mean ± SEM) from three independent experiments with a total of 6 western blot analyses (See Fig. S4).

### RNA immunoprecipitation and qRT-PCR (RIP-qRT-PCR)

Bacterial RNA immunoprecipitation was performed as described previously (52). Briefly, strains IU1945 (WT) and IU5809 (*cvfD*-L-FLAG^3^) were grown in 400 mL of BHI broth to OD_620_ of 0.25 to 0.40. Cells were collected by centrifugation (8,000 X *g* for 10 min at 4°C). Cell pellets were washed once with 30 mL of cold 1 X PBS (4°C) and resuspended in 2 mL of cold lysis buffer (50 mM Tris-HCl pH 7.4, 150 mM NaCl, 1% Triton X100 (v/v) containing protease inhibitor (ThermoFisher; cat# 78429). Suspensions (2 mL) were transferred into two lysing matrix B tubes (1 mL each) (MP Biomedicals, Inc.). Tubes were shaken in a FastPrep homogenizer at 4°C for ten 40s cycles (4X shaking; 5 min on ice; 3X shaking; 5 min on ice; 3X shaking) at a setting of 6.0 M/s. Cell debris was removed by centrifugation at 16,000 X *g* for 5 min at 4°C. Protein concentration was determined by DC™ protein assay (Bio-Rad). About 1 mL of lysate with similar protein concentration was added to tubes with 50 μL of anti-FLAG magnetic beads (Sigma) and rotated for 2 h at 4°C. The beads were washed thrice with 1 mL of lysis buffer for 10 min with rotation at 4°C. FLAG-tagged protein bound to RNA was eluted by incubation with 100 μL of FLAG elution solution (150 ng of 3X FLAG peptide per μL) (Sigma) for 30 min at 4°C. RNA was extracted by phenol-chloroform extraction. Briefly, 100 μL of phenol/chloroform/isoamyl alcohol (125:24:1, pH 4.3, ThermoFisher) was added to the eluate and vortexed for 10 sec before incubation at room temperature for 5 min. The mixture was centrifuged at 16,000 X *g* for 5 min. The upper phase was transferred to a new 1.5 mL tube. 2 volumes of isopropanol and 1/10 volume of 3 M sodium acetate (pH 5.2) was added to the aqueous phase, followed by incubation for 10 min at −80°C to precipitate RNA. Precipitated material was collected by centrifugation (16,000 X *g* for 30 min at 4°C) and washed with 500 μL of 80% ethanol. Pellets were dried at room temperature for 10 min and suspended in 30 μL of nuclease-free water. 5 μg of purified RNA was treated by DNase from a DNA-free DNA removal kit (Ambion). Reverse transcription and qRT-PCR were performed as described above. Ratio of *phoU2* transcript abundance between CvfD-L-FLAG^3^ and untagged WT samples was used as enrichment ratio. 16s rRNA was used as control.

### Mouse models of infection

All procedures were approved by the Bloomington Institutional Animal Care and Use Committee (BIACUC) and were performed according to recommendations of the National Research Council. Experiments were performed as described in (91) with the following changes. Male ICR mice (21-24 g; Harlan) were anaesthetized by inhaling 4% isoflurane (Butler animal Health Supply) for 8 min. In two independent experiments, a total of 8 mice were intranasally inoculated with each specific bacterial strain to be tested. Bacteria were grown exponentially in BHI to OD_620_ of ~0.1. Ten milliliters of culture were centrifuged for 5 min at 14,500 X *g* and then suspended in 1 mL 1 X PBS to yield ~10^7^ CFU mL^−1^. CFU counts were confirmed by serial dilution and plating. Fifty microliters of suspensions were administered as described previously (72). Mice were monitored every 8 h intervals and moribund mice were euthanized by CO_2_ asphyxiation, which was used as the time of death in statistical analyses. Kaplan-Meir survival curves and log-rank tests were generated using GraphPad Prism 5 software.

## SUPPLEMENTAL MATERIALS

Supplemental Materials are available for this article.

## ACKNOWLEDGEMENTS

We thank Kurt Zimmer and Doug Rusch for assistance with Illumina DNA sequencing and mRNA-Seq analyses. This work was supported by the McGovern Medical Startup funds (to D.S. and N.R.D.) and NIGMS grants RO1GM127715, RO1GM128439, and R35GM131767 (M.E.W).

## Notes

### Competing Interest Statement

The authors have declared no competing interest.

