## Supplemental Tables and Figures for "S1-Domain RNA-Binding Protein (CvfD) Is a New Post-Transcriptional Regulator That Mediates Cold Sensitivity, Phosphate Transport, and Virulence in *Streptococcus pneumoniae* D39"

**Running Title:** Pleiotropic phenotypes of pneumococcal Δ*cvfD* mutants

**Keywords:** RNA-binding protein; post-transcriptional regulation; phosphate uptake; cold sensitivity

† Dhriti Sinha and Jiaqi J. Zheng contributed equally to this work, and author order was determined both alphabetically and in order of increasing seniority.

#### TABLE OF CONTENTS

**SUPPLEMENTAL TABLES**

**TABLE S1.** *S. pneumoniae* strains used in this study

**TABLE S2.** Primers used to construct mutants used in this study

**TABLE S3.** Growth characteristics of strains used in this study

**TABLE S4.** Mutations in Δ*cvfD* suppressor mutants determined by Illumina whole-genome sequencing

**TABLE S5.** Oligonucleotide primers used for qRT-PCR

**SUPPLEMENTAL FIGURES**

**FIGURE S1.** Encapsulated D39 Δ*cvfD* (IU4772) showed decreased growth yield and increased doubling time with increased time interval between the preparation and the usage of the BHI media

**FIGURE S2.** Encapsulated D39 Δ*cvfD* (IU4772) strain shows drastically reduced growth compared to its *cvfD*⁺ parent when incubated at 32°C on TSAII BA plates

**FIGURE S3** qRT-PCR analysis to validate the changes in the relative transcript levels of *trpD*, *spd_0104* and *spd_1874* as identified by RNA-seq

**FIGURE S4.** Representative anti-HA Western blots to determine the half-life of PhoU2-HA in *cvfD*⁺ and Δ*cvfD* strains

**FIGURE S5.** Growth characteristics of the Δ*cvfD* suppressor mutants

**FIGURE S6.** Defective growth of Δ*cvfD* (IU4772) mutant in a chemically defined medium lacking added $Mn^{2+}$ (CDM-Mn) can be improved by the addition of 500 μM of $Mn^{2+}$

**SUPPLEMENTAL REFERENCES**

**Table S1.** *S. pneumoniae* strains used in this study

| Strain | Genotype (description)[a] | Antibiotic resistance[b] | Reference or source |
|---|---|---|---|
| K243 | D39 Δ*cps* Δ*spd_1366*::$P_c$-[*kan*$^R$-*rpsL*$^+$] (IU1945 transformed with Δ*spd_1366*::$P_c$-[*kan*$^R$-*rpsL*$^+$] amplicon) | Kan$^R$ | This study |
| E253 | D39 Δ*cps* Δ*spd_1366*::$P_c$-*erm* (IU1945 transformed with Δ*spd_1366*::$P_c$- *erm* amplicon) | Erm$^R$ | This study |
| IU1690 | D39W | None | NCTC 7466; Lanie et al., 2007 (1); Slager et al., 2018 (1a) |
| IU1781 | D39 *rpsL1* | Str$^R$ | Ramos-Montanez et al., 2008 (2) |
| IU1945 | D39 Δ*cps* | None | Lanie et al., 2007 (1) |
| IU2173 | D39 *rpsL1* Δ*spxB* | Str$^R$ | Ramos-Montanez et al., 2008 (2) |
| IU3116 | D39 *rpsL1* CEP::$P_c$-[*kan*$^R$-*rpsL*$^+$] | Kan$^R$ | Ramos-Montanez et al., 2010 (3) |
| IU3286 | D39 *rpsL1* Δ*cps2E::*$P_c$-[*kan*$^R$-*rpsL*$^+$] | Kan$^R$ | Ramos-Montanez et al., 2010 (3) |
| IU3309 | D39 *rpsL1 cps2E* (ΔA) | Str$^R$ | Ramos-Montanez et al., 2010 (3) |
| IU4707 | D39 *rpsL1* Δ*spd_1366*::$P_c$-[*kan*$^R$-*rpsL*$^+$] (IU1781 transformed with Δ*spd_1366*::$P_c$-[*kan*$^R$-*rpsL*$^+$] amplicon from K243) | Kan$^R$ | This study |
| IU4708 | D39 *rpsL1* Δ*spd_1366*::$P_c$-[*kan*$^R$-*rpsL*$^+$] (IU1781 transformed with Δ*spd_1366*::$P_c$-[*kan*$^R$-*rpsL*$^+$] amplicon from K243; different isolate) | Kan$^R$ | This study |
| IU4771 | D39 *rpsL1* Δ*spd_1366* (IU4707 transformed with Δ*spd_1366* amplicon) | Str$^R$ | This study |
| IU4772 | D39 *rpsL1* Δ*spd_1366* (IU4708 transformed with Δ*spd_1366* amplicon) | Str$^R$ | This study |
| IU5506 | D39 *rpsL1* Δ*spd_1366* CEP::$P_c$-[*kan*$^R$-*rpsL*$^+$] (IU4772 transformed with CEP::$P_c$-[*kan*$^R$-*rpsL*$^+$] amplicon from IU3116) | Kan$^R$ | This study |
| IU5508 | D39 *rpsL1* Δ*spd_1366 bgaA::Kan-T1T2-*$P_{ftsA}$-*spd_1366* (IU4772 transformed with *bgaA::Kan-T1T2-*$P_{ftsA}$-*spd_1366* amplicon) | Str$^R$ Kan$^R$ | This study |
| IU5809 | D39 Δ*cps spd_1366*-L-FLAG$^3$ -$P_c$-*erm* | Erm$^R$ | Zheng et al., 2017 (4) |

| | | | |
|---|---|---|---|
| IU6139 | D39 *rpsL1* Δ*phoU2*::$P_c$-[*kan*$^R$-*rpsL*$^+$] | Kan$^R$ | Zheng et al., 2016 (5) |
| IU7291[c] | D39 *rpsL1* Δ*spd_1366*$^{\text{Suppressor isolate1}}$ | Str$^R$ | This study |
| IU7293[d] | D39 *rpsL1* Δ*spd_1366*::$P_c$-[*kan*$^R$-*rpsL*$^+$]$^{\text{Suppressor isolate2}}$ | Str$^R$ | This study |
| IU7294[e] | D39 *rpsL1* Δ*spd_1366*$^{\text{Suppressor isolate3}}$ | Str$^R$ | This study |
| IU8269 | D39 *rpsL1* Δ*spd_1366* Δ*cps2E::*$P_c$-[*kan*$^R$-*rpsL*$^+$] (IU4772 transformed with Δ*cps2E::*$P_c$-[*kan*$^R$-*rpsL*$^+$] amplicon from IU3286) | Kan$^R$ | This study |
| IU8396 | D39 *rpsL1* Δ*spd_1366* *cps2E* (ΔA) (IU8269 transformed with *cps2E* (ΔA) amplicon from IU3309) | Str$^R$ | This study |
| IU8675 | D39 *rpsL1* *phoU2*-HA (IU6139 transformed with *phoU2*-HA amplicon) | Str$^R$ | This study |
| IU8717 | D39 *rpsL1* *cvfD*-L-FLAG$^3$-$P_c$-*erm* (IU1781 transformed with *cvfD*-L-FLAG$^3$-$P_c$-*erm* amplicon from IU5809) | Str$^R$ Erm$^R$ | This study |
| IU8719 | D39 *rpsL1* *phoU2*-HA Δ*spd_1366*:: $P_c$-*erm* *bgaA::Kan-T1T2-*$P_{ftsA}$*-spd_1366* (IU8722 transformed with *bgaA::Kan-T1T2-*$P_{ftsA}$*-spd_1366* amplicon from IU5508) | Str$^R$ Erm$^R$ Kan$^R$ | This study |
| IU8722 | D39 *rpsL1* *phoU2*-HA Δ*spd_1366*:: $P_c$-*erm* (IU8675 transformed with Δ*spd_1366*:: $P_c$-*erm* amplicon from E253) | Str$^R$ Erm$^R$ | This study |
| IU17714 | D39 *rpsL1* Δ*spxB* Δ*spd_1366*::$P_c$-[*kan*$^R$-*rpsL*$^+$] (IU2173 transformed with Δ*spd_1366*::$P_c$-[*kan*$^R$-*rpsL*$^+$] amplicon from K243) | Kan$^R$ | This study |
| NRD10125[f] | D39 *rpsL1* Δ*spd_1366*::$P_c$-[*kan*$^R$-*rpsL*$^+$] (IU1781 transformed with Δ*spd_1366*::$P_c$-[*kan*$^R$-*rpsL*$^+$]; independent amplicon generated using fusion PCR) | Kan$^R$ | This study |
| NRD10133[g] | D39 *rpsL1* Δ*spd_1366* (NRD10125 transformed with Δ*spd_1366* amplicon; independent amplicon generated using fusion PCR) ) | Str$^R$ | This study |

[a] Strains were constructed by transformation of amplicons into the indicated recipient strain as described in Materials and Methods. Primers used to synthesize fusion amplicons are listed in Table S2. FLAG and HA (Hemagglutinin) epitope tags were introduced by creating carboxyl terminal translational fusions. The amino acid sequences of FLAG and HA are DYKDDDDK and YPYDVPDYA respectively.

[b] Antibiotic resistance markers: Kan, kanamycin; Str, streptomycin; Erm, Erythromycin.

[c] Isolated as a spontaneous growth suppressor of IU4771 during growth at 37$^oC$ in BHI.

[d] Isolated as a spontaneous growth suppressor of IU4708 during growth at 37$^oC$ in BHI.

[e] Isolated as a spontaneous growth suppressor of IU4772 during growth at 37°C in BHI in the presence of 200 μM Zn(II).

[f, g] Strains used in sRNA-seq.

[f] Strain has the same genotype as IU4707 and IU4708; created via a separate transformation reaction using an independently created PCR amplicon (Table S2).

[g] Strain has the same genotype as IU4771 and IU4772; created via a separate transformation reaction using an independently created PCR amplicon (Table S2).

**Table S2.** Primers used to construct mutants used in this study

| Primer name | Primer Sequence (5'-3') | Template[a] | Product |
|---|---|---|---|
| **For construction of strain K243 (Δ*spd_1366*::$P_c$-[*kan*$^R$-*rpsL*$^+$])** | | | |
| P594 | AAGGGAGATGACTTGCACTTGCCTGACAGT | D39 | Upstream of *spd_1366* + 60 bp of 5' *spd_1366* |
| P596 | CATTATCCATTAAAAATCAAACGGATCCTAAAGGCACCGTAGGGCTGAATC | | |
| Kan rpsL forward | TAGGATCCGTTTGATTTTTAATGGATAATG | $P_c$-[*kan-rpsL*$^+$] cassette[b] | $P_c$-[*kan-rpsL*$^+$] |
| Kan rpsL reverse | GGGCCCCTTTCCTTATGCTTTTG | | |
| P597 | CAAAAGCATAAGGAAAGGGGCCCCGACGAATGCTACCAATCTGGACT | D39 | 60 bp of 3' *spd_1366* + downstream |
| P595 | ACTGGCAAGCGTGCTCGTGTATATTTGGCT | | |
| **For construction of strain E253 (Δ*spd_1366*::$P_c$-*erm*)** | | | |
| P594 | AAGGGAGATGACTTGCACTTGCCTGACAGT | D39 | Upstream of *spd_1366* + 60 bp of 5' *spd_1366* |
| P596 | CATTATCCATTAAAAATCAAACGGATCCTAAAGGCACCGTAGGGCTGAATC | | |
| Kan rpsL forward | TAGGATCCGTTTGATTTTTAATGGATAATG | $P_c$-*erm* cassette[b] | $P_c$-*erm* |
| Kan rpsL reverse | GGGCCCCTTTCCTTATGCTTTTG | | |
| P597 | CAAAAGCATAAGGAAAGGGGCCCCGACGAATGCTACCAATCTGGACT | D39 | 60 bp of 3' *spd_1366* + downstream |
| P595 | ACTGGCAAGCGTGCTCGTGTATATTTGGCT | | |
| **For construction of strain IU4771, IU4772 (Δ*spd_1366*)** | | | |
| P594 | AAGGGAGATGACTTGCACTTGCCTGACAGT | D39 | Upstream of *spd_1366* + 60 bp of 5' *spd_1366* |
| DM117 | GCTTCTCCAGTCCAGATTGGTAGCATTCGTCGAAAGGCACCGTAGGGCTGAATCCCT | | |
| DM116 | GTATTACAGGGATTCAGCCCTACGGTGCCTTTCGACGAATGCTACCAATCTGGACTG | D39 | 60 bp of 3' *spd_1366* + downstream |
| P595 | ACTGGCAAGCGTGCTCGTGTATATTTGGCT | | |
| **For construction of strain IU5508 (*bgaA::Kan-T1T2-P$_{ftsA}$-spd_1366*)** | | | |
| P146 | TGGCCATTCATCGCTGGTCGTGCTGAAAT | IU4981[c] | *bgaA::Kan-T1T2-P$_{ftsA}$* |
| DM146 | CTTAGTCCTCGATTTCAATAGTTTCAATACATCGCTTCCTCTCTATCTTCCAAG | | |
| DM145 | GGAAGATAGAGAGGAAGCGATGTATTGAAACTATTGAAATCGAGGACTAAGATG | D39 | *spd_1366* |
| DM187 | CAACTGGTTTATGAGAAAGTAAGTTCTTCTACTTCTTCTTTTTTAAATGGTGCAGGG | | |
| DM186 | TGCACCATTTAAAAAAGAAGAAGTAGAAGAACTTACTTTCTCATAAACCAGTTGC | IU4981[c] | 3' *bgaA*' |
| CS121 | GCTTTCTTGAGGCAATTCACTTGGTGC | | |
| **For construction of strain IU8675 (phoU2-HA)** | | | |
| P1257 | TGTATCGCTCGTGCCATCTCTGTTAAGCCA | D39 | *phoU2*-HA |

| | | | |
|---|---|---|---|
| JQ107 | AGAATTAAACTATTTTAAGCATAATCTGGAACATCAT ATGGATATAGTTCGACAATCTTA | | |
| JQ108 | TAAGATTGTCGAACTATATCCATATGATGTTCCAGAT TATGCTTAAAATAGTTTAATTCT | D39 | HA + downstream of *phoU2* |
| P1258 | CAGCCTGCAATTCATTGACTGCTTCACCCA | | |
| **For construction of strain NRD10125 (Δ*spd_1366*::$P_c$-[*kan*$^R$-*rpsL*$^+$])** | | | |
| Spd_1366 For | GGACTTTTGAAGATAAGGGAGATG | D39 | Upstream of *spd_1366* + 60 bp of 5’ *spd_1366* |
| Spd_1366 FusRev | CATTATCCATTAAAAATCAAACGGATCCTAAAAGGCA CCGTAGGGCTGAATCCCTGTAATACG | | |
| Kan rpsL forward | TAGGATCCGTTTGATTTTTAATGGATAATG | $P_c$-[*kan-rpsL*$^+$] cassette[b] | $P_c$-[*kan-rpsL*$^+$] |
| Kan rpsL reverse | GGGCCCCTTTCCTTATGCTTTTG | | |
| Spd_1366 FusFor | CGACGAATGCTACCAATCTGGACTGGAGAAGCCCT GCACCATTTAAAAAAGAAGAAGTAG | D39 | 60 bp of 3' *spd_1366* + downstream |
| Spd_1366 Rev | GCTTCTTCCTTACTGGCAAAACC | | |
| **For construction of strain NRD10133 (Δ*spd_1366*)** | | | |
| Spd_1366 For | GGACTTTTGAAGATAAGGGAGATG | D39 | Upstream of *spd_1366* + 60 bp of 5’ *spd_1366* |
| Spd_1366 cleanRev | CAGTCCAGATTGGTAGCATTCGTCGAAAGGCACCGT AGGGCTGAATCCCTGTAATAC | | |
| Spd_1366 cleanFor | GTATTACAGGGATTCAGCCCTACGGTGCCTTTCGAC GAATGCTACCAATCTGGACTG | D39 | 60 bp of 3' *spd_1366* + downstream |
| Spd_1366 Rev | GCTTCTTCCTTACTGGCAAAACC | | |

[a] Genomic DNA of indicated *S. pneumoniae* strains was used as templates for PCR reactions, except for $P_c$-[*kan-rpsL*$^+$] and $P_c$-*erm* cassettes.

[b] $P_c$-*erm* and $P_c$-[*kan-rpsL*$^+$] cassettes are described in (Tsui *et al.*, 2011) (6).

[c] IU4981 is an unpublished strain with genotype *bgaA::Kan-T1T2*-$P_{ftsA}$-*divIVA.* $P_{ftsA}$ is the intergenic DNA sequence upstream of *ftsA* (4).

**Table S3.** Growth characteristics of strains used in this study

| Strain | Genotype | Doubling time (min)[a,c] | Growth yield ($OD_{620}$)[b,c] | n[d] |
|---|---|---|---|---|
| **Growth in BHI at 37$^{O}$C** | | | | |
| IU1781 | WT (D39 *rpsL1*) | 44.8 ± 0.8 | 0.89 ± 0.01 | 14 |
| NRD10133 | Δ*cvfD* | 53.2 ± 1.3** | 0.35 ± 0.02** | 3 |
| IU4772 | Δ*cvfD* | 61.5 ± 2.5 *** | 0.26 ± 0.02*** | 11 |
| IU5508 | Δ*cvfD bgaA::Kan-T1T2-cvfD*$^{+}$ | 43.2 ± 2.0$^{ns}$ | 0.78 ± 0.04** | 3 |
| IU7291 | Δ*cvfD*$^{\text{Suppressor isolate1}}$ | 45.8 ± 0.6$^{ns}$ | 0.71 ± 0.02** | 3 |
| IU7293 | Δ*cvfD*::$P_c$-[*kan*$^{R}$-*rpsL*$^{+}$]$^{\text{Suppressor isolate2}}$ | 45.3 ± 0.5$^{ns}$ | 0.51 ± 0.07** | 3 |
| IU7294 | Δ*cvfD*$^{\text{Suppressor isolate3}}$ | 46.5 ± 3.1$^{ns}$ | 0.5 ± 0.06** | 3 |
| IU8675 | *phoU2*-HA | 43.7 ± 2.2$^{ns}$ | 0.81 ± 0.04* | 3 |
| IU8722 | *phoU2*-HA Δ*cvfD* | 56.5 ± 6.8** | 0.30 ± 0.09*** | 3 |
| **Growth in BHI at 37$^{O}$C** | | | | |
| IU3309 | *cps2E* (ΔA) | 37.5 ± 0.6 | 0.99 ± 0.01 | 2 |
| IU8396 | Δ*cvfD cps2E* (ΔA) | 42.2 ± 0.6** | 0.77 ± 0.05* | 5 |
| **Growth in BHI at 32$^{O}$C** | | | | |
| IU1781 | WT | 57.9 ± 1.2 | 0.84 ± 0.01 | 5 |
| IU4772 | Δ*cvfD* | 109.2 ± 9.6*** | 0.08 ± 0.01*** | 5 |
| IU5508 | Δ*cvfD bgaA::Kan-T1T2-cvfD* | 63.3 ± 9.5$^{ns}$ | 0.78 ± 0.03$^{ns}$ | 3 |
| **Growth in BHI at 32$^{O}$C** | | | | |
| IU3309 | *cps2E* (ΔA) | 53.9 ± 2.1 | 0.94 ± 0.01 | 2 |
| IU8396 | Δ*cvfD cps2E* (ΔA) | 57.1 ± 2.3** | 0.79 ± 0.03* | 5 |
| **WT growth in BHI broth containing Mn and/or Zn at 37$^{O}$C[e]** | | | | |
| IU1781 | WT | 45.5 ± 1.5 | 0.88 ± 0.02 | 6 |
| IU1781 + Zn(II) | WT | 56.2 ± 2.6** | 0.76 ± 0.02*** | 6 |
| IU1781 + Mn(II) | WT | 42.5 ± 1.6$^{ns}$ | 0.97 ± 0.01* | 4 |
| IU1781 + Zn(II) + Mn(II) | WT | 42.0 ± 2.9$^{ns}$ | 0.96 ± 0.00$^{ns}$ | 2 |
| **IU4772 growth in BHI broth containing Mn and/or Zn at 37$^{O}$C[e]** | | | | |
| IU4772 | Δ*cvfD* | 65.2 ± 3.8 | 0.24 ± 0.02 | 6 |
| IU4772 + Zn(II) | Δ*cvfD* | 66.5 ± 2.3$^{ns}$ | 0.14 ± 0.02** | 6 |
| IU4772 + Mn(II) | Δ*cvfD* | 47.4 ± 2.7** | 0.59 ± 0.02*** | 4 |
| IU4772 + Zn(II) + Mn(II) | Δ*cvfD* | 46.9 ± 2.9* | 0.65 ± 0.00*** | 2 |
| **WT growth in CDM-Mn medium containing Mn and/or Zn at 37$^{O}$C[e]** | | | | |
| IU1781 | WT | 75.8 ± 5.1 | 1.17 ± 0.02 | 2 |
| IU1781 + Mn(II) | WT | 60.3 ± 0.3$^{ns}$ | 1.27 ± 0.04$^{ns}$ | 2 |
| IU1781 + Zn(II) + Mn(II) | WT | 62.6 ± 0.6$^{ns}$ | 1.29 ± 0.06$^{ns}$ | 2 |

| | | | | |
|---|---|---|---|---|
| **IU4772 growth in CDM-Mn medium containing Mn and/or Zn at 37$^{O}$C[e]** | | | | |
| IU4772 | Δ*cvfD* | 145.0 ± 7.5 | 0.68 ± 0.08 | 2 |
| IU4772 + Mn(II) | Δ*cvfD* | 82.1 ± 1.8* | 1.06 ± 0.06$^{ns}$ | 2 |
| IU4772 + Zn(II) + Mn(II) | Δ*cvfD* | 91.8 ± 4.1* | 1.07 ± 0.08$^{ns}$ | 2 |

[a] Doubling times were determined from data points that showed exponential growth as shown with a semi-log plot of $OD_{620}$ vs time.

[b] Maximum growth yield is the highest $OD_{620}$ value obtained during a 9 to 10 h, or 12 h period after subculture to a starting inoculum of $OD_{620}$ ~ 0.005 in BHI, or CDM-Mn, respectively.

[c] Mean values of doubling times and growth yields are presented with ± SEM and are calculated using GraphPad Prism version 7. ns, *,**,*** indicate that the values are nonsignificant, or p values are < 0.05, 0.01 or 0.001, respectively, when compared with the strain in the first line of each table using an unpaired, two-tailed t-test with GraphPad Prism 7.

[d] n indicates the number of independent replicates.

[e] Growth of WT or IU4772 in BHI broth or CDM-Mn medium containing added $Mn^{2+}$ and $Zn^{2+}$: +Mn(II), 0.5 mM $MnSO_4$; +Zn(II) + Mn(II), 0.2 mM $ZnCl_2$ + 0.5 mM $MnSO_4$.

**Table S4.** Mutations in Δ*cvfD* suppressor mutants determined by Illumina whole-genome sequencing

| Strain | Gene containing indicated mutation | Function | Amino acid change |
|---|---|---|---|
| **IU7291** ($\Delta cvfD^{\text{Suppressor isolate1}}$) | | | |
| | 754 bp deletion including ~193 bp of 3′ region of *spxB,* entire *spd_0637* and 103 bp downstream of *spd_0637* | SpxB, pyruvate oxidase; SPD_0637, glyoxalase family protein | |
| | *spd*_2003 (*dltC*) (GAA → TAA) | D-alanyl carrier protein subunit involved in incorporation of D-Ala into membrane-associated D-alanyl lipoteichoic acid | Glu9Stop |
| | *spd*_0851 (*pyrK*) (CAG → CGG) | Dihydroorotate dehydrogenase involved in pyrimidine biosynthesis | Gln104Arg |
| **IU7293** ($\Delta cvfD^{\text{Suppressor isolate2}}$) | | | |
| | *spd*_0319 (*cps2E*) (CGT → AGT) | Undecaprenylphosphate glucosephosphotransferase | Arg307Ser |
| | *spd*_1170 (GGC → TGC) | Oligopeptide ABC transporter, oligopeptide binding protein | Thr463Cys |
| **IU7294** ($\Delta cvfD^{\text{Suppressor isolate3}}$) | | | |
| | *spd*_0581 (*thyA*) (GCC → ACC) | Thymidylate synthase, involved in pyrimidine biosynthesis | Ala155Thr |
| | *spd*_1719 (TCG → TAG) | PAP2 family protein | Ser98Stop |
| | *spd*_0792 (GTC → GGC) | Lipoprotein, putative | Val229Gly |

**Table S5.** Oligonucleotide primers used for qRT-PCR

| Primer name | Primer Sequence (5'-3') | Gene |
|---|---|---|
| KW096 | CCAAACAGTCAGCTTCAGGAACGA | *pstS1* |
| KW097 | AGATCCCAAGGAGATGTAGCCGAT | |
| JQ43 | AGCCACAAGTGTCTGACCTTCGAT | *phoU1* |
| JQ44 | TTGAGGCTTGGTGCAAAGGAAAGG | |
| JQ45 | ATCGCACTCCAACAACCAGTCTCT | *phoU2* |
| JQ46 | GATCAAGTGCTGCTTCAACAACGC | |
| AL031 | CTGCGACAGCAGATTTGACCACTA | *spd_1874* |
| AL032 | TTCCTGAGGAGCTTCTTCTGCAAC | |
| AL023 | GGAGTAGCTGCCTTGTTTGCAGTA | *spd_0104* |
| AL024 | CAGGGCCATCGATAACCAACTCTT | |
| JQ101 | CTGTTATCCTGATGGCGATTGTTC | *czcD* |
| JQ102 | CCAAAGCATCCATAGTCCAGAGAT | |
| JQ105 | AGTCAACTGTCTGGAGGTCAATTC | *psaA* |
| JQ106 | CTTGATCGAAGTAGTGGGGAATCT | |
| KK461 | CTGGCTTGAACGGAACAACCAAGA | *trpD* |
| KK462 | CCAGCATTCAAGACTGTCGTTTCC | |
| DM270 | GTATTACAGGGATTCAGCCCTAC | *spd_1366* (*cvfD*) |
| DM271 | CTCCAAAGTGCGGATAGAAAGA | |
| KK387 | AAAGGTCGTGGTGGTAAGGGAATG | *gyrA* |
| KK388 | GCATCTTGATCCAGGCGCATTACT | |
| KK489 | CAGCAGTAGGGAATCTTCGGCAAT | *16S rRNA* |
| KK490 | TACGCCCAATAAATCCGGACAACG | |

#### SUPPLEMENTAL FIGURES

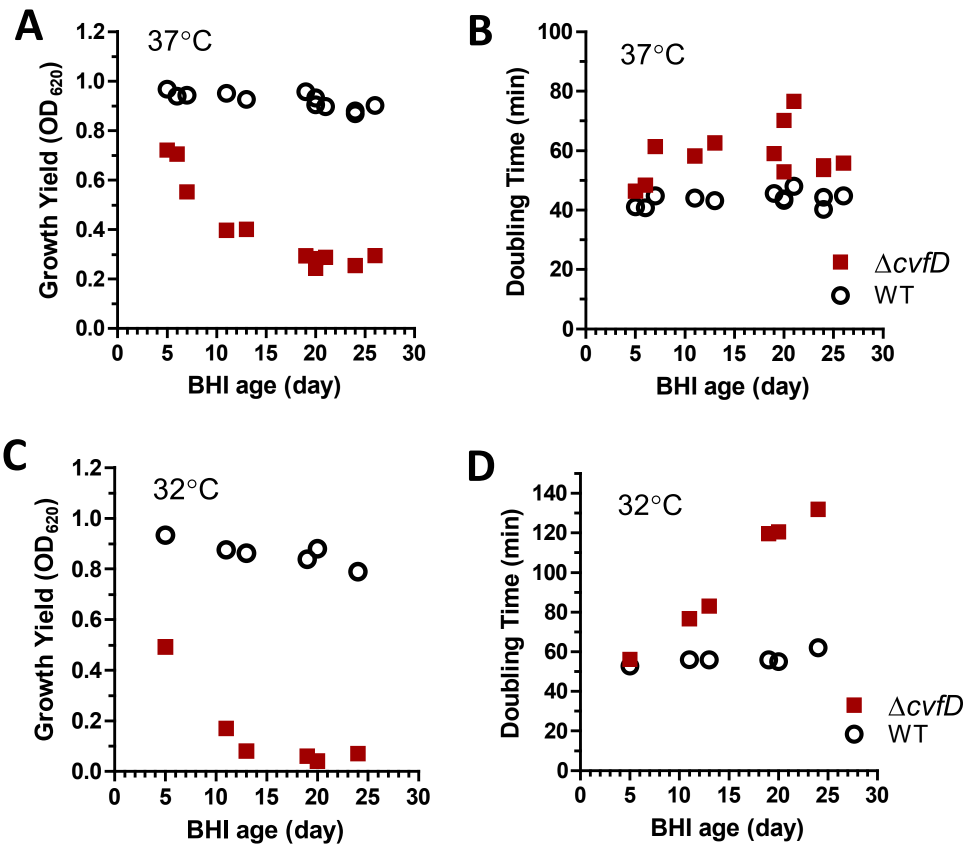


**Figure S1. Encapsulated D39 Δ*cvfD* (IU4772) showed decreased growth yield and increased doubling time with increased time interval between the preparation and the usage of the BHI media.** Growth yields (A, C) and doubling times (B, D) of encapsulated D39 parent strain (IU1781) and its derived mutant strain Δ*cvfD* (IU4772) in BHI broth that were autoclaved, and stored for a period of time before use for growth experiment at 37°C (A and B) or 32°C (C and D). Each data point represents the result from individual independent growth experiment for which the date of BHI broth was recorded. Table S3 includes data from BHI that were 19 days or older from this set of experiments and other growth data that generate similar values for the Δ*cvfD* strain.

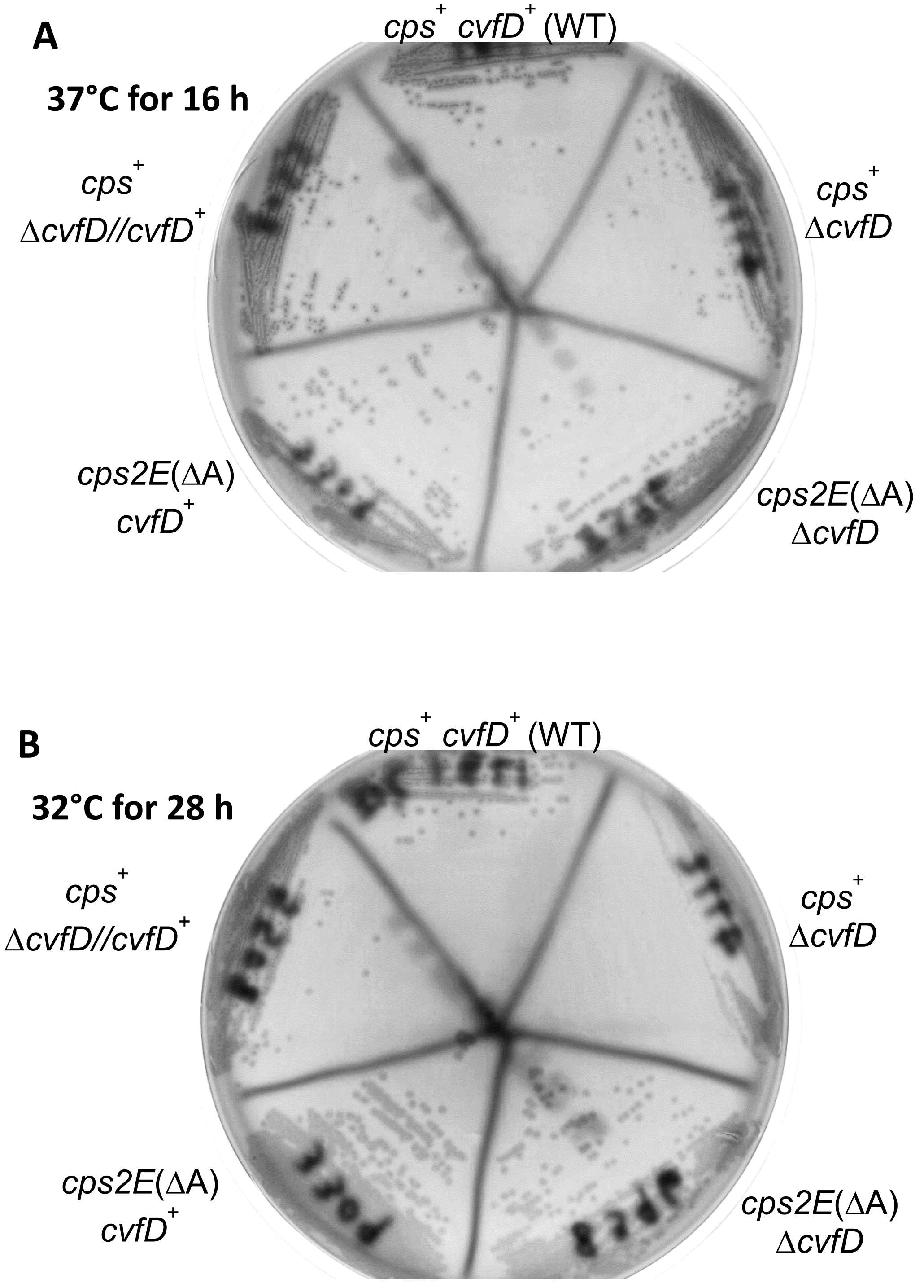


**Figure S2. Encapsulated D39 Δ*cvfD* (IU4772) strain shows drastically reduced growth compared to its *cvfD*$^+$ parent when incubated at 32°C on TSAII BA plates.** Strains IU1781 (*cps*$^+$ *cvfD*$^+$ WT), IU4772 (*cps*$^+$ Δ*cvfD*), IU5508 (*cps*$^+$ Δ*cvfD*//*cvfD*$^+$), IU3309 (*cps2E*(ΔA) *cvfD*$^+$) and

IU8396 (*cps2E*(ΔA) Δ*cvfD*) were first streaked from frozen ice stock onto TSAII BA plates and incubated at 37°C overnight. Single colonies were then streaked onto fresh TSAII BA plates and placed in a 37°C (A) or in a 32°C (B) incubator. After incubation for 16 h at 37°C (A), the colony sizes of *csp*$^+$ Δ*cvfD* strain were slightly smaller than the *cps*$^+$ *cvfD*$^+$ parent strain or the complemented *cps*$^+$ Δ*cvfD*//*cvfD*$^+$ strain. In contrast, after incubation for 28 h at 32°C, healthy colonies were present with *cps*$^+$ *cvfD*$^+$ parent strain or the complemented *cps*$^+$ Δ*cvfD*//*cvfD*$^+$ strain, but no single colonies were visible with the *csp*$^+$ Δ*cvfD* strain.  Growth of *cvfD*$^+$ and Δ*cvfD* in the unencapsulated *cps2E*(ΔA) genetic background were similar to each other at both 37°C and 32°C.

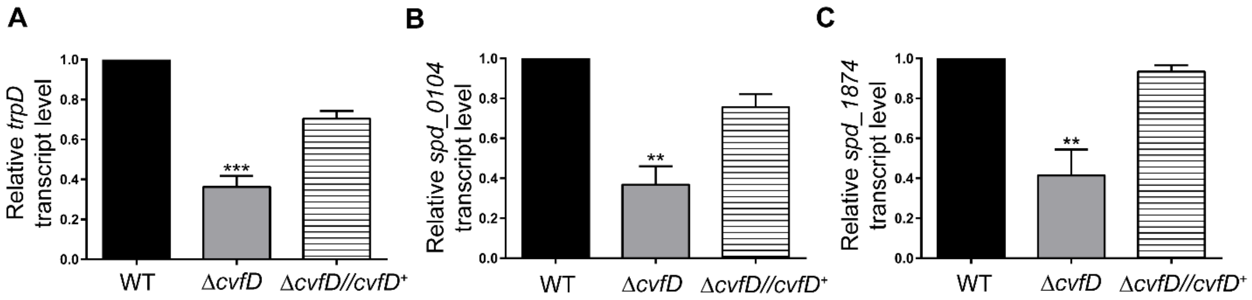


**Figure. S3. qRT-PCR analysis to validate the changes in the relative transcript levels of *trpD*, *spd_0104* and *spd_1874* as identified by RNA-seq.** *trpD* (A), *spd_0104* (B) and *spd_1874* (C) transcript levels were determined in a wild-type D39 parent (WT; IU1781) and its derived isogenic mutants (Δ*cvfD*; IU4772 and Δ*cvfD bgaA*::$P_{ftsA}$ *cvfD*$^{+}$; IU5508) as described previously in the legend of Fig. 3. Data points and error bars in the graphs represent the mean (± SEM) of at least three independent experiments.  qRT-PCR primers used are listed in Supplemental Table S5.

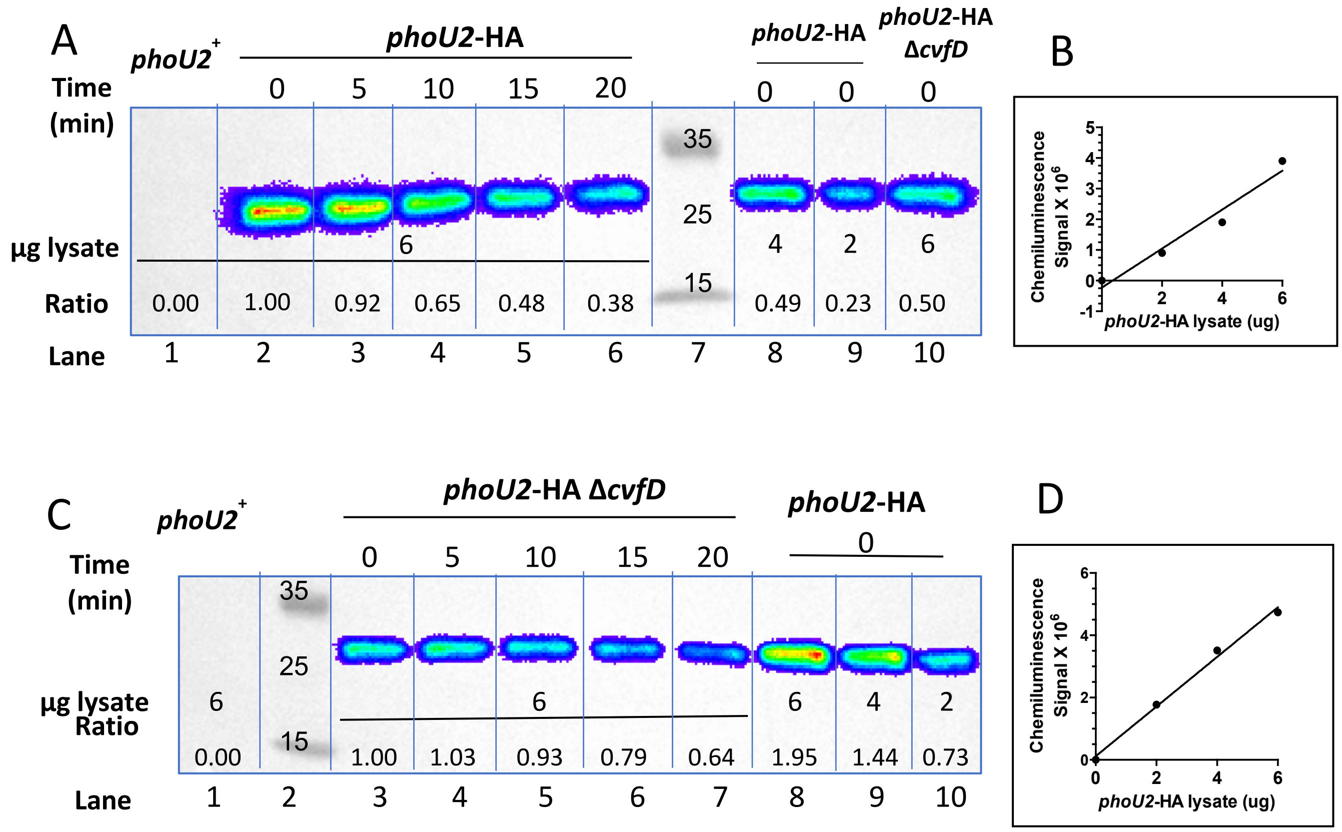


**Figure S4. Representative anti-HA Western blots to determine the half-life of PhoU2-HA in *cvfD*$^+$ and Δ*cvfD* strains.** (A) A representative anti-HA Western blot to determine the half-life of PhoU2-HA of IU8675 (*phoU2*-HA). Strains used for lysates preparation, and time (min) of treatment with Cm are indicated above the lanes. Amounts of lysate (µg protein) loaded per lane are indicated below the bands. Ratio indicates the relative amount of PhoU2-HA signal relative to *phoU2*-HA strain at $T_0$ of Cm treatment (lane 2) as extrapolated from a standard curve shown in (B), which was generated using signal intensities obtained from lanes loaded with 2 µg (lane 9), 4 µg (lane 8), or 6 µg (lane 2) of *phoU2*-HA lysates. The standard curve demonstrates that the chemiluminescent signals obtained with the Cm-treated samples (lanes 2 to 6) were in the linear range of Western detection. The ratio of PhoU2-HA in *phoU2*-HA Δ*cvfD* (lane 10) at $T_0$ relative to that in *phoU2*-HA *cvfD*$^+$ strain at $T_0$ (lane 2) is 0.5 from this Western analysis. (C) A representative anti-HA Western blot to determine the half-life of PhoU2-HA of IU8722 (*phoU2*-HA Δ*cvfD*). Ratio indicates the relative amount of PhoU2-HA signal relative to *phoU2*-HA Δ*cvfD* strain at $T_0$ of Cm treatment (lane 3) as extrapolated from a standard curve shown in (D), which was generated using signals obtained from lanes loaded with 2 μg (lane 10), 4 μg (lane 9), or 6 μg (lane 8) of lysates obtained from *phoU2*-HA strain. The ratio of PhoU2-HA in *phoU2*-HA $cvfD^+$ (lane 8) at $T_0$ relative to that in *phoU2*-HA Δ*cvfD* strain at $T_0$ (lane 3) is 1.95 from this Western analysis. Expected molecular weight of PhoU2-HA is 26.2 kDa.

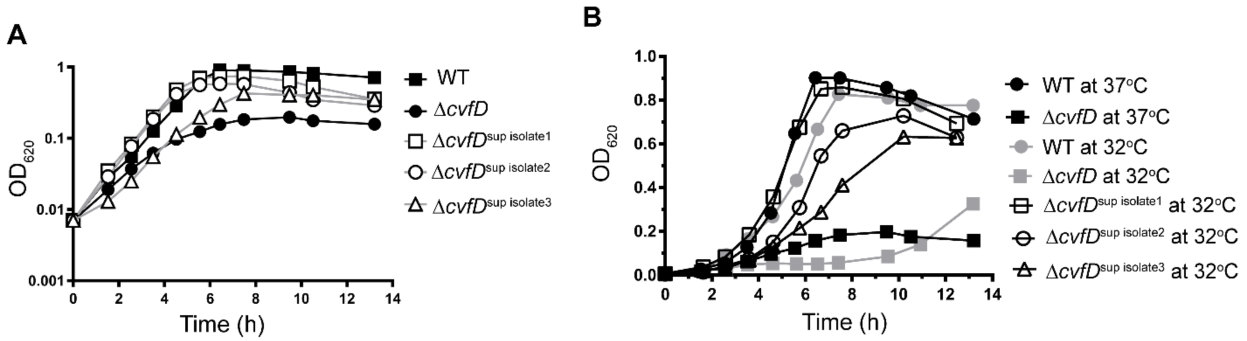


**Figure S5. Growth characteristics of the Δ*cvfD* suppressor mutants.** Representative growth curves of the encapsulated D39 parent strain (IU1781) and its derived mutant strains Δ*cvfD* (IU4772), Δ*cvfD*$^{supressor\ isolate1}$ (IU7291), Δ*cvfD*$^{supressor\ isolate2}$ (IU7293) and Δ*cvfD*$^{supressor\ isolate3}$ (IU7294) at 37 °C (A) and at 32 °C (B). At least three independent growth curves were performed yielding similar results, and a representative curve is shown.  Average growth rates and growth yields are listed in Supplemental Table S3.

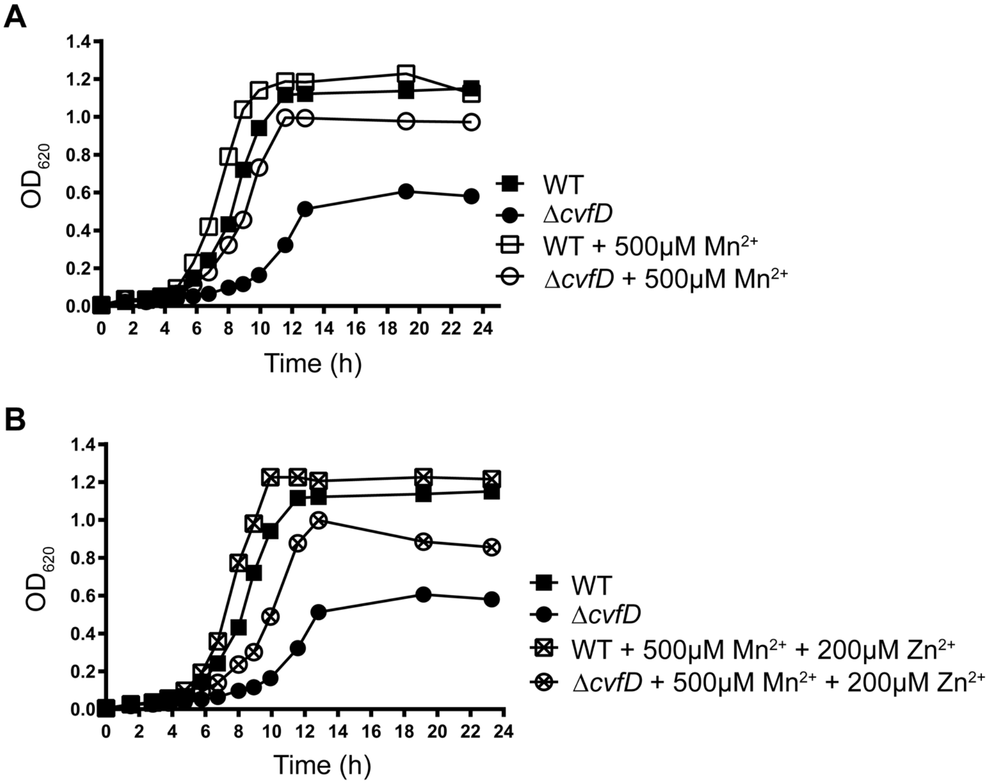


**Figure S6: Defective growth of $\Delta$*cvfD* (IU4772) mutant in a chemically defined medium lacking added $Mn^{2+}$ (CDM-Mn) can be improved by the addition of 500 μM of $Mn^{2+}$.** (A) Growth characteristics of $\Delta$*cvfD* (IU4772) mutant in CDM-Mn in the presence or absence of 500 μM of $Mn^{2+}$ relative to the wild-type parent (WT; IU1781) at 37 $^{o}$C. (B) Representative growth curves of a $\Delta$*cvfD* mutant (IU4772) relative to parent (IU1781) in CDM-Mn in the presence of both 500 μM of $Mn^{2+}$ and 200 μM $Zn^{2+}$. (A, B) At least two independent growth curves were performed under the above-mentioned conditions yielding similar results, and a representative curve is shown. Average growth rates and growth yields are listed in Supplemental Table S3.
